# Humanized TfR1 and transferrin gene-replacement rats for in vivo evaluation of BBB transport

**DOI:** 10.1101/2025.07.25.666792

**Authors:** Metin Yesiltepe, Sanjay Metkar, Tao Yin, Ishita Chakraborty, Luciano D’Adamio

## Abstract

The transferrin receptor 1 (TfR1)–transferrin (TF) axis plays a central role in iron homeostasis and has long been recognized as a promising route for delivering biologics across the blood–brain barrier (BBB). We have developed a class of human-specific anti-TfR1 nanobodies (NewroBus) that exploit this transport pathway. However, the lack of cross-reactivity with rodent TfR1 limits the utility of standard animal models for preclinical testing. To overcome this challenge, we generated knock-in (KI) rats in which the coding sequences of the endogenous *Tfrc* and *Tf* genes were replaced with human coding sequences, yielding animals that express human TfR1 and/or human TF under physiological control. Rats homozygous for both humanized alleles were viable and fertile, indicating that the human proteins can functionally replace their rodent equivalents. Nonetheless, these double homozygous rats exhibited erythropoietic abnormalities and tissue-specific alterations in iron distribution—characterized by decreased splenic and increased hepatic iron—suggesting incomplete functional compensation. In contrast, heterozygous rats showed only mild, subclinical hematologic changes (microcytosis and hypochromia). These findings demonstrate that the humanized TfR1–TF axis is compatible with life and iron regulation, albeit with varying degrees of compensation depending on gene dosage. Importantly, these KI rats provide a translationally relevant platform for evaluating pharmacokinetics, CNS penetration, and safety of human-specific BBB-targeting therapeutics, including NewroBus-based biologics and other TfR1-mediated delivery strategies.

## Introduction

Delivering biologics to the central nervous system (CNS) is a challenge due to the limited permeability of the blood-brain barrier (BBB). One popular strategy to overcome this obstacle is receptor-mediated transcytosis via transferrin receptor 1 (TfR1), which enables transport of cargo across the BBB (1–6).

We developed nanobodies against TfR1—referred to as NewroBus (see reference (7) and accompanying paper)—because they are small, stable, easy to engineer, and suitable for fusion to therapeutic payloads. Their compact size facilitates efficient BBB transcytosis, and they can be produced at high yield using cost-effective systems. As monomers, they are less likely to induce internalization of plasma membranes TfR1 homodimers. NewroBus nanobodies have been shown to efficiently mediate the transport of otherwise BBB-impermeable anti-TNFα inhibitory nanobodies (8) across the BBB in vivo (see reference (7) and accompanying paper).

To reduce the risk of immune responses in clinical applications, NewroBus nanobodies were humanized using *in silico* AI/machine learning-guided deimmunization tools. Humanized variants were validated through *ex vivo* immunogenicity assays with human immune cells and functional testing to confirm retained activity. These studies showed very low—if any—immunogenic potential.

NewroBus nanobodies were selected to avoid interfering with the transferrin (Tf)–TfR1 interaction, which is essential for iron uptake and metabolism (see Ref. (7) and accompanying paper). Tf binds iron (Fe³⁺) tightly at neutral pH and delivers it into cells by binding to TfR1 or TfR2, triggering endocytosis. Inside the acidic endosome, iron is released, and the Tf–TfR complex is recycled back to the cell surface, where Tf detaches (9). Disrupting this process can be toxic—particularly by causing anemia—because TfR1 is vital for iron delivery to cells, red blood cell formation, and nervous system development. Mice lacking both copies of the *Tfrc* gene (*Tfrc* knockout) die during embryonic development due to severe anemia and developmental defects. Mice with one copy (heterozygotes) are viable but show signs of iron deficiency, such as smaller and less pigmented red blood cells (10). Additionally, mice with very low transferrin levels also develop severe anemia (11).

Functionally important regions of TfR1—such as the helical and protease-like domains involved in Tf binding and internalization—are highly conserved across species (>85–90% identity between human and rodent TfR1; ∼95–98% with monkey TfR1). In contrast, surface-exposed loops not involved in these functions are much less conserved (<60% identity with rodents; ∼75–85% with monkeys). Because of the selection strategy of avoiding key functional interference, NewroBus nanobodies most likely bind to non-functional, surface-exposed loops of TfR1 that are structurally accessible but not critical for receptor function. This hypothesis is currently being tested by CryoEM structural analysis of the NewroBus–TfR1 complex. This targeting strategy explains why NewroBus nanobodies lack of cross-species reactivity and only bind human TfR1 and not rodent or monkey TfR1 (see Ref. (7) and accompanying paper), because it targets poorly conserved regions.

While this human specificity improves safety, it limits the use of conventional animal models. Accurate *in vivo* studies require humanized models expressing human TfR1 to assess NewroBus function. To address this, we generated TfR1 knock-in rats using a gene replacement strategy that preserves physiological expression. In parallel, to evaluate potential toxicity related to transferrin binding, we also generated rats expressing human transferrin. Here, we characterize these humanized rat models and evaluate their utility for testing the brain delivery and safety of NewroBus nanobody carriers.

## Results and Discussion

### Generation of humanized *TFRC* and *TF* Knock-in rats

To enable preclinical testing of human-specific NewroBus-based therapeutics, we generated gene-replacement rat models expressing human TfR1. In parallel, we developed rats expressing human transferrin to better assess potential toxicity arising from interference with human Tf–TfR1 interactions. Due to the large size of both the rat and human genes, we used a targeted strategy in which the human coding sequences were inserted into exon 2 of the endogenous rat *Tf* and *Tfrc* loci. In the resulting KI alleles, the human TfR1 protein is expressed as a full-length type II membrane protein, while the TF protein retains the rat signal peptide fused to the human TF secreted domain. During translation, this rat leader sequence is cleaved, producing fully human TF protein. This strategy was chosen to favor a physiologically regulated expression of the human proteins under the control of the endogenous rat promoters and regulatory elements (**Figure 1**).

**Figure 1.**
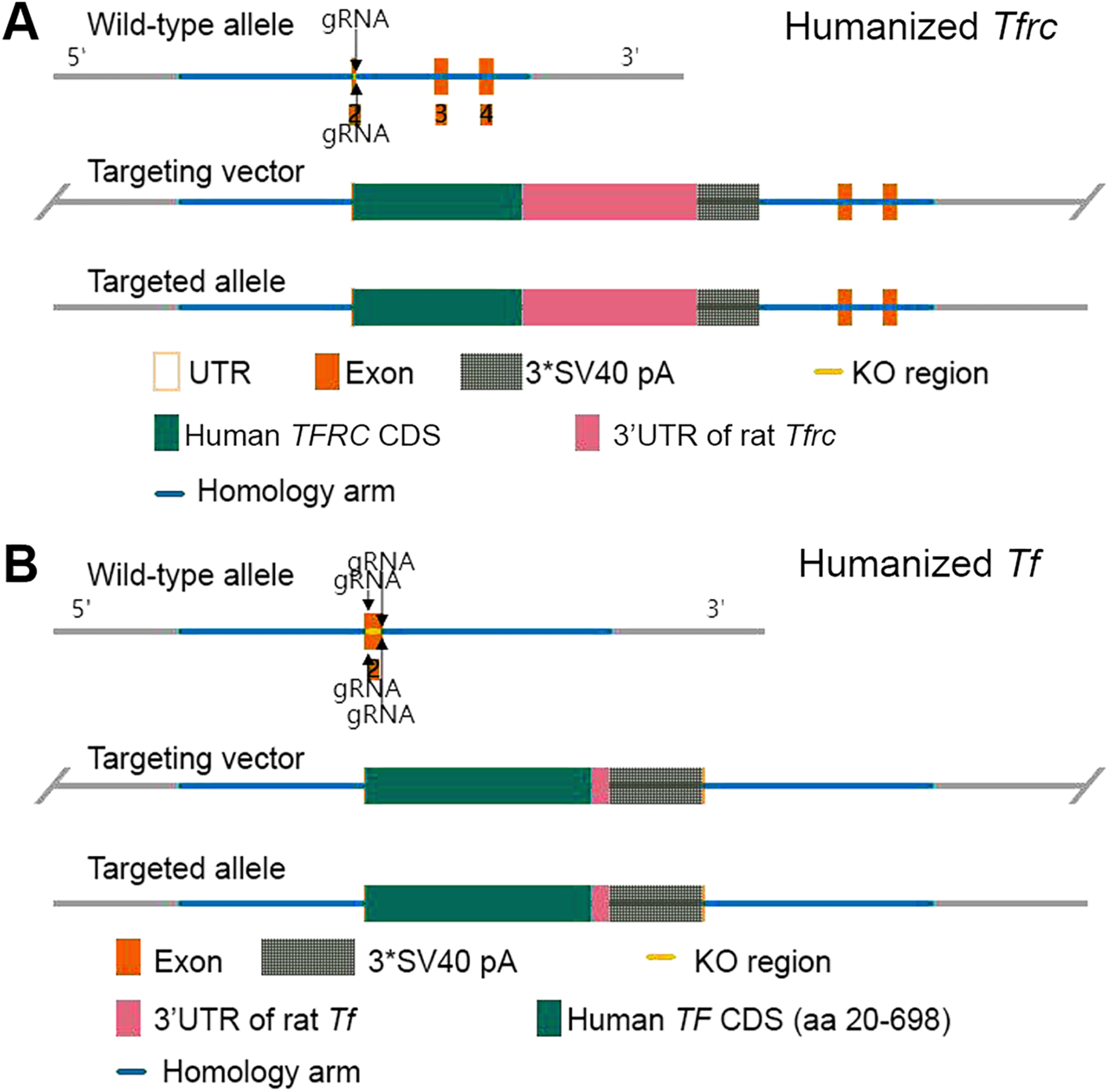
Generation of humanized *Tf and Tfrc* rats. **A.** The rat *Tfrc* gene consists of 19 exons, with the ATG start codon located in exon 2 and the TAA stop codon in exon 19. In our KI model, the last 32 base pairs (3’) of rat *Tfrc* gene exon 2, counting from the ATG start codon, were replaced with the human *TFRC* CDS (Transcript: 201-ENST00000360110), followed by the rat 3’ untranslated region (3’UTR) and the SV40 polyadenylation sequence. **B**. The rat *Tf* gene (NM_001013110.1) is comprised of 17 exons, with the ATG start codon located in exon 1 and the TAA stop codon in exon 17. In the KI model, a portion of exon 2 coding sequence (140 base pairs) of the rat *Tf* gene was substituted with the Human *TF* CDS (amino acids 20-698, Transcript: 201-ENST00000402696), followed by the rat 3’UTR and the SV40 polyadenylation sequence. Importantly, the murine signal peptide (amino acids 1-19), encoded by exon 1 and the 5’ sequences of rat exon 2, was retained. Consequently, this modified allele results in a TF protein encompassing the rat leader sequence and the human TF protein.

### Evaluation of viability and postnatal growth in KI rat models

We crossed double heterozygous rats carrying one humanized and one endogenous allele at each locus—*Tf^h/w^:Tfrc^h/w^*—to assess whether specific combinations of these gene replacements result in early lethality or other phenotypes suggestive of deleterious systemic effects and developmental incompatibility. In this nomenclature, “^h^” denotes the human knock-in allele and “^w^” refers to the wild-type rat allele.

Genotype frequencies followed Mendelian ratios, with no significant deviation detected by Chi-square analysis comparing observed versus expected genotype counts (χ² = 2.281, df = 8, p = 0.9711), indicating no embryonic lethality associated with humanized *Tf* or *Tfrc* KI alleles (**Figure 2A**). Survival analysis at weaning (about 24 days of age) showed no significant differences between genotypes or sexes. Chi-square analysis comparing survival versus death across genotypes revealed no significant association (χ² = 8.167, df = 8, p = 0.4173), suggesting that heterozygous or homozygous expression of human *TF* and *TFRC* does not affect early postnatal viability (**Figure 2B**).

**Figure 2.**
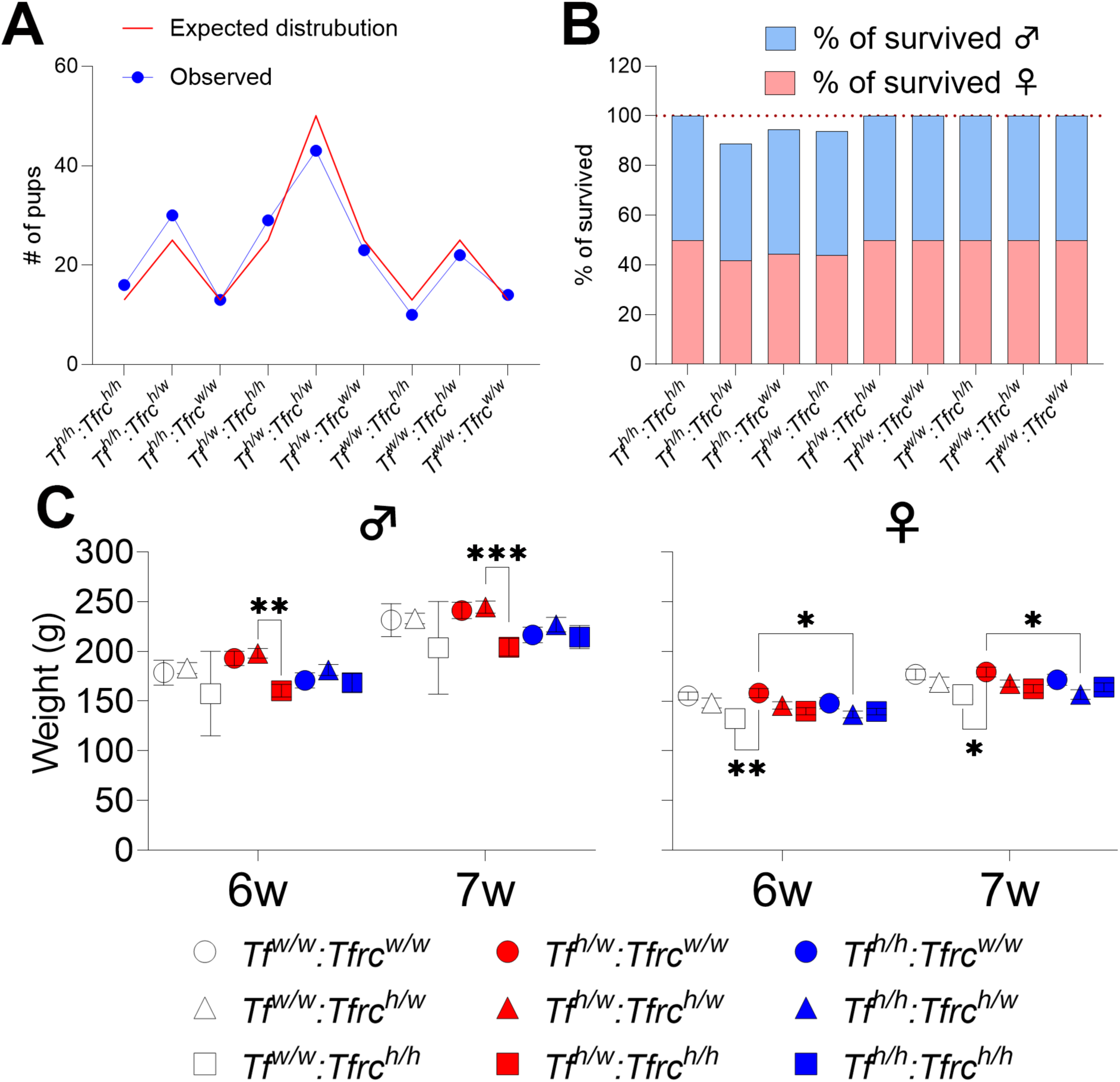
Survival and growth in humanized KI rats. **A.** Genotype distribution of offspring from heterozygous intercrosses showed no significant deviation from expected Mendelian ratios, indicating no embryonic lethality associated with the humanized alleles. **B.** Survival analysis at weaning revealed no significant genotype- or sex-dependent differences in early postnatal viability. **C**. Body weights of rats across genotypes and sexes. No significant genotype-dependent differences were observed in body weight, indicating that TF and TfR1 humanization do not affect overall growth under baseline conditions. Data are presented as mean ± SEM. Statistical comparisons were performed using appropriate post hoc tests; refer to text for detailed p-values.

To assess whether the humanized *Tf* and *Tfrc* KI alleles affect general health or development, we measured body weights in male and female rats at 6 and 7 weeks of age. Animals were grouped by genotype: *Tf^w/w^:Tfrc^w/w^*, *Tf^w/w^:Tfrc^h/w^*, *Tf^w/w^:Tfrc^h/h^*, *Tf^h/w^:Tfrc^w/w^*, *Tf^h/w^:Tfrc^h/w^*, *Tf^h/w^:Tfrc^h/h^*, *Tf^h/h^:Tfrc^w/w^*, *Tf^h/h^:Tfrc^h/w^*, *Tf^h/h^:Tfrc^h/h^*. Measurements were analyzed separately by sex to account for sex-dependent growth differences. All genotypes showed a normal increase in body weight from week 6 to week 7 (**Figure 2C**). However, genotype-specific differences emerged. Statistical comparisons revealed that *Tf^h/w^:Tfrc^h/h^* males weighed significantly less than *Tf^h/w^:Tfrc^h/w^* at both 6 weeks (p=0.006) and 7 weeks (p=0.0007). In females, *Tf^w/w^:Tfrc^h/h^* animals weighed significantly less than *Tf^h/w^:Tfrc^w/w^* at both time points (week 6, p=0.006; week 7, p=0.0297). Similarly, *Tf^h/h^:Tfrc^h/w^* females also weighed significantly less than *Tf^h/w^:Tfrc^w/w^* at both 6 weeks (p=0.0273) and 7 weeks (p=0.0144).

These findings demonstrate that the humanized *Tf* and *Tfrc* alleles are inherited in a Mendelian fashion and do not impair postnatal survival or general growth. In mice, *Tfrc* knockout causes embryonic lethality, and *Tf* deficiency leads to perinatal death unless rescued by transfusion (11). In contrast, all humanized rats— including double homozygous *Tfrc* and *Tf* knock-in animals—are viable and fertile. This indicates that the human *Tfrc* and *Tf* alleles can functionally replace their rat counterparts, assuming these genes serve similar essential roles in rats as they do in mice.

### Expression analysis of human *TF* and *TFRC* mRNA

To assess whether the humanized *Tf* and *Tfrc* knock-in alleles retain physiological expression patterns, we compared human and endogenous rat mRNA levels in double heterozygous *Tf^h/w^:Tfrc^h/w^* rats. qRT-PCR was used to measure transcript levels across five organs (brain, heart, kidney, liver, spleen) from eight animals (n=4 per sex). Because the TaqMan assays target exon–exon junctions and the knock-in alleles are intron-less—mimicking mature mRNA—we ran PCR on each RNA sample without reverse transcription. The –RT controls showed no amplification (Figure S1b), confirming the absence of genomic DNA that could otherwise yield false-positive signals.

*TF* is primarily synthesized in the liver in both humans and rodents (12); accordingly and as expected, both human *TF* and rat *Tf* were predominantly expressed in the liver and barely detectable in other tissues (**Figure 3**). Hepatic expression levels of both human *TF* (2way ANOVA summary F_(1,30)_= 7.233 p=0.0116, for liver female vs. male p<0.0001) and rat *Tf* (2way ANOVA summary F_(1,30)_= 4.594 p=0.0403, for liver female vs. male p=0.0002) were higher in males compared to females, indicating a sex-dependent difference in liver transferrin expression for both the wild-type and humanized alleles.

**Figure 3.**
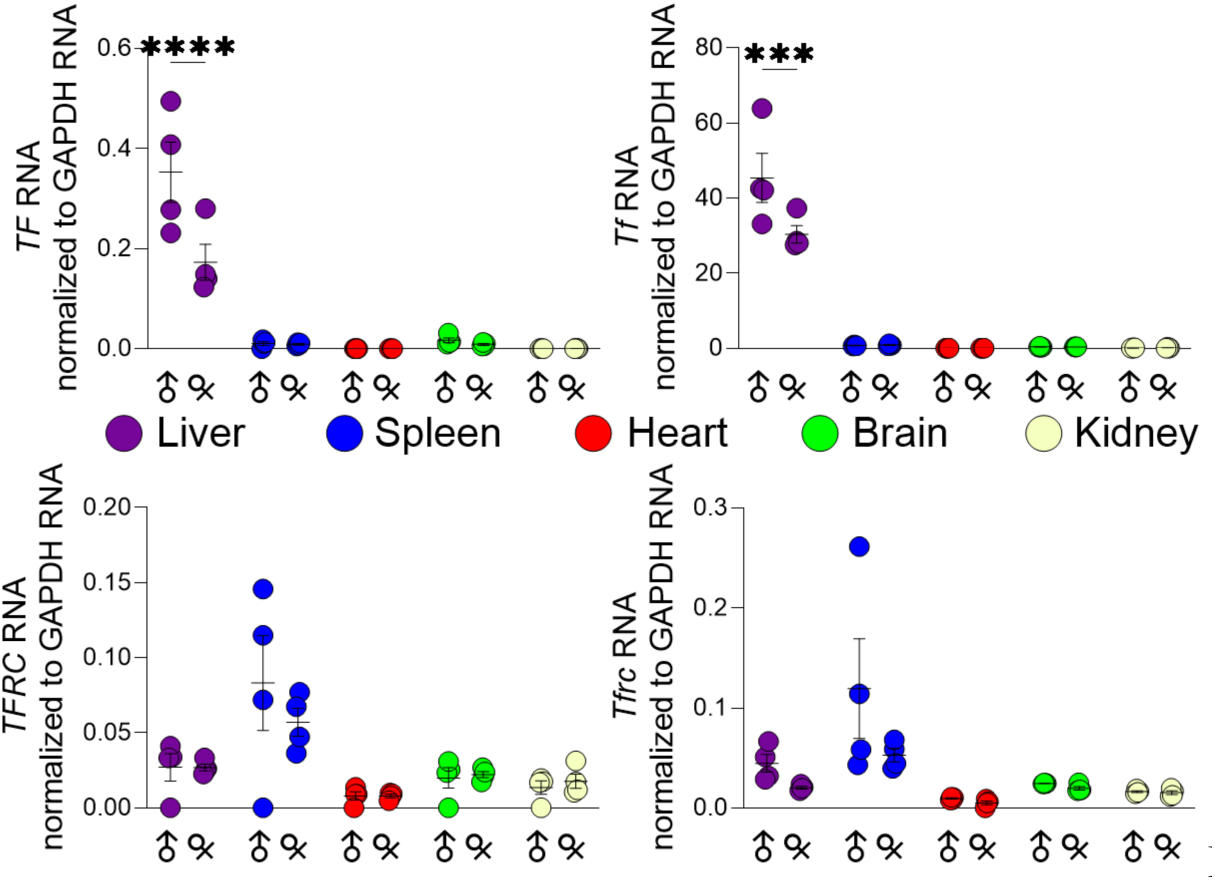
Tissue-Specific mRNA and protein expression of TF and TfR1 in heterozygous KI rats. qRT-PCR analysis of human *TFRC* and *TF* (left panels) and rat *Tfrc* and *Tf* (right panels) was performed on brain, heart, kidney, liver, and spleen tissues from male and female *Tf^h/w^*:*Tfrc^h/w^* rats (n = 4 per sex). Human *TF* expression was largely restricted to the liver, similar to rat *Tf*, which was also liver-enriched (upper panels). Human *TFRC* mRNA was most abundant in the spleen, mirroring the expression pattern of endogenous rat *Tfrc* (lower panels). No significant sex differences were observed in *TFRC* expression, while both human and rat *TF* expressions were higher in males than in females in the liver.

Human *TFRC* expression was highest in the spleen, followed by liver and kidney. Rat *Tfrc* showed a similar tissue distribution, with comparable expression levels across these organs. No significant sex-dependent differences were observed for either transcript (2way ANOVA summary; for human *TFRC* F_(1,30)_= 0.3079 p=0.5831, for rat *Tfrc* F_(1,30)_= 3.915 p=0.0571) (**Figure 3**). These findings suggest that the humanized *Tf* and *Tfrc* alleles are transcriptionally regulated in a tissue- and sex-specific manner consistent with their endogenous rat counterparts.

### Detection of human TF and sTfR1 in CSF and serum

As previously noted, TF is a secreted glycoprotein primarily produced by the liver and released into the bloodstream, where it functions as the principal iron transport protein (13). In contrast, TfR1 is a membrane-bound receptor expressed on the cell surface that mediates cellular iron uptake through binding and internalization of TF. A portion of surface TfR1 can be cleaved by metalloproteases, generating a soluble form (sTfR1) that circulates in the blood and serves as a biomarker of iron status and erythropoietic activity (14).

To assess whether the humanized TF and TfR1 alleles in our humanized rat models support proper expression, processing, and secretion of the human proteins, we measured human TF and sTfR1 concentrations in both CSF and serum. All nine genotype combinations—wild-type, heterozygous, and homozygous for both *Tf* and *Tfrc*—were included to evaluate the effects of gene dosage on protein expression and secretion.

In homozygous *Tf^h/h^* rats, serum human TF levels ranged from 200 to 550 µg/mL (**Figure 4C**). Serum TF levels in heterozygous *Tf^h/w^* rats were approximately half of those observed in homozygous *Tf^h/h^* animals, consistent with gene-dosage-dependent expression of the humanized allele. In healthy humans, the physiological serum range for TF is 1,750–3,600 µg/mL (15). indicating that human TF expression in the KI rats is markedly lower than in humans. This discrepancy may be due to species-specific differences in protein processing, including altered clearance kinetics resulting from the lower binding affinity of human TF to rat TfR1, reduced hepatic expression of the human TF gene in rat hepatocytes, or suppression of TF production as a negative acute-phase reactant. Serum TF levels were significantly higher in genotypes co-expressing human TfR1 (**Figure 4C**; all statistical comparisons in Table 1), suggesting that the presence of human TfR1— which has a higher affinity for human TF than the rat receptor—may enhance the stability and/or reduce the clearance of circulating human TF *in vivo*. Human TF was also detected in the CSF of knock-in rats in a gene-dosage–dependent manner, but at concentrations approximately 200-fold lower than those measured in serum (**Figure 4A**). Due to the lack of reliable published reference values for TF concentrations in CSF, direct comparisons to normal physiological levels in either species cannot be made at this time.

**Figure 4.**
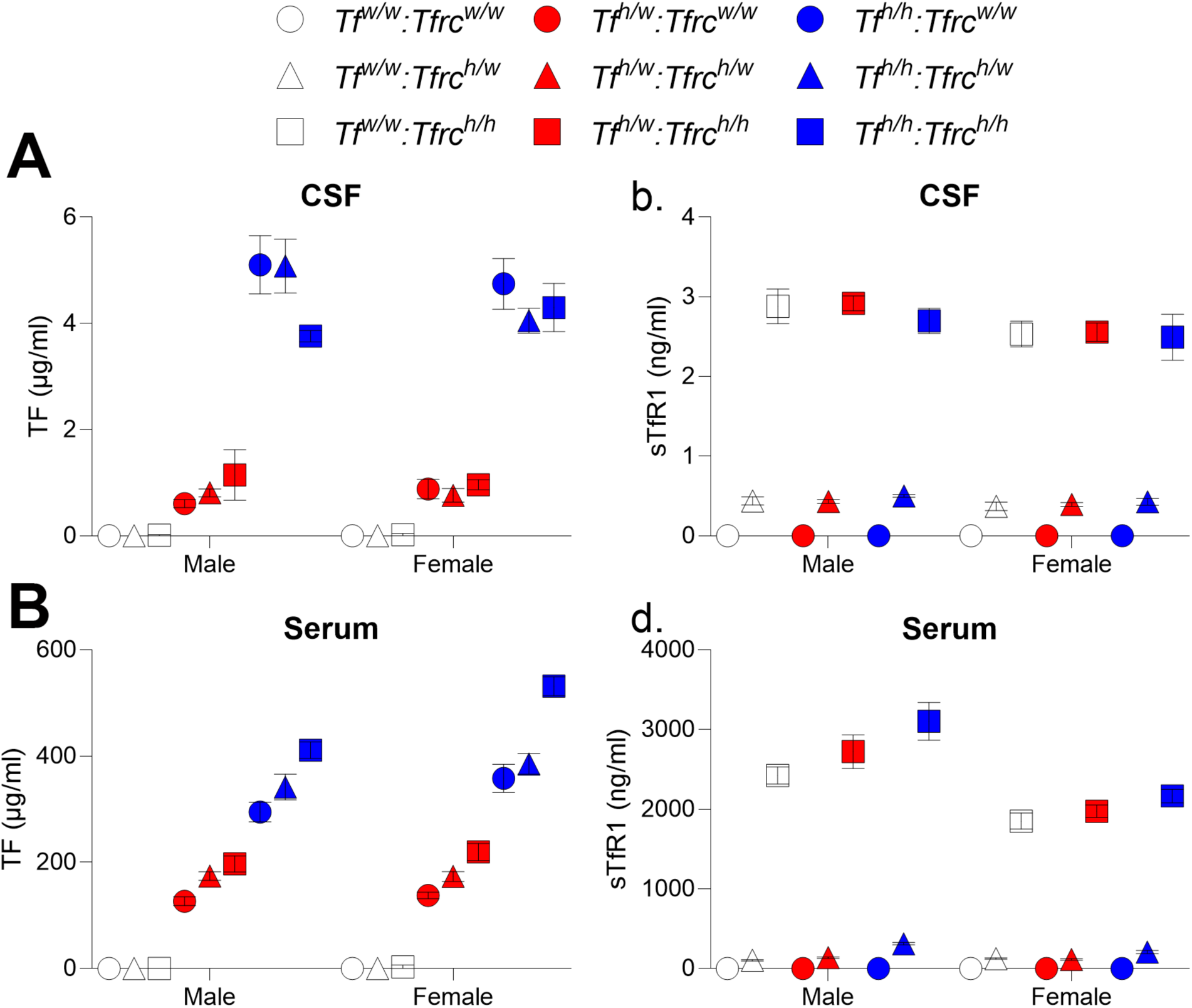
Detection of human Transferrin (TF) and soluble TfR1 (sTfR1) in serum and CSF of KI rats. Human-specific ELISA assays were used to measure concentrations of TF and sTfR1 in serum and CSF from rats carrying various combinations of wild-type and humanized *TF* and *TFRC* alleles. **(A,B)** Human TF was detected in both serum and CSF of *Tf^h/w^* and *Tf^h/h^* animals, with highest levels observed in *Tf^h/h^* genotypes. **(C,D)** Human sTfR1 was present only in animals carrying humanized *TFRC* alleles, with levels corresponding to gene dosage: *Tfrc^h/h^* > *Tfrc^h/w^*; undetectable in *Tfrc^w/w^* rats. Data represent mean ± SEM. Statistical comparisons were performed using two-way ANOVA with Tukey’s post hoc test; exact p-values are reported in **Table 1**.

**Table 1.**
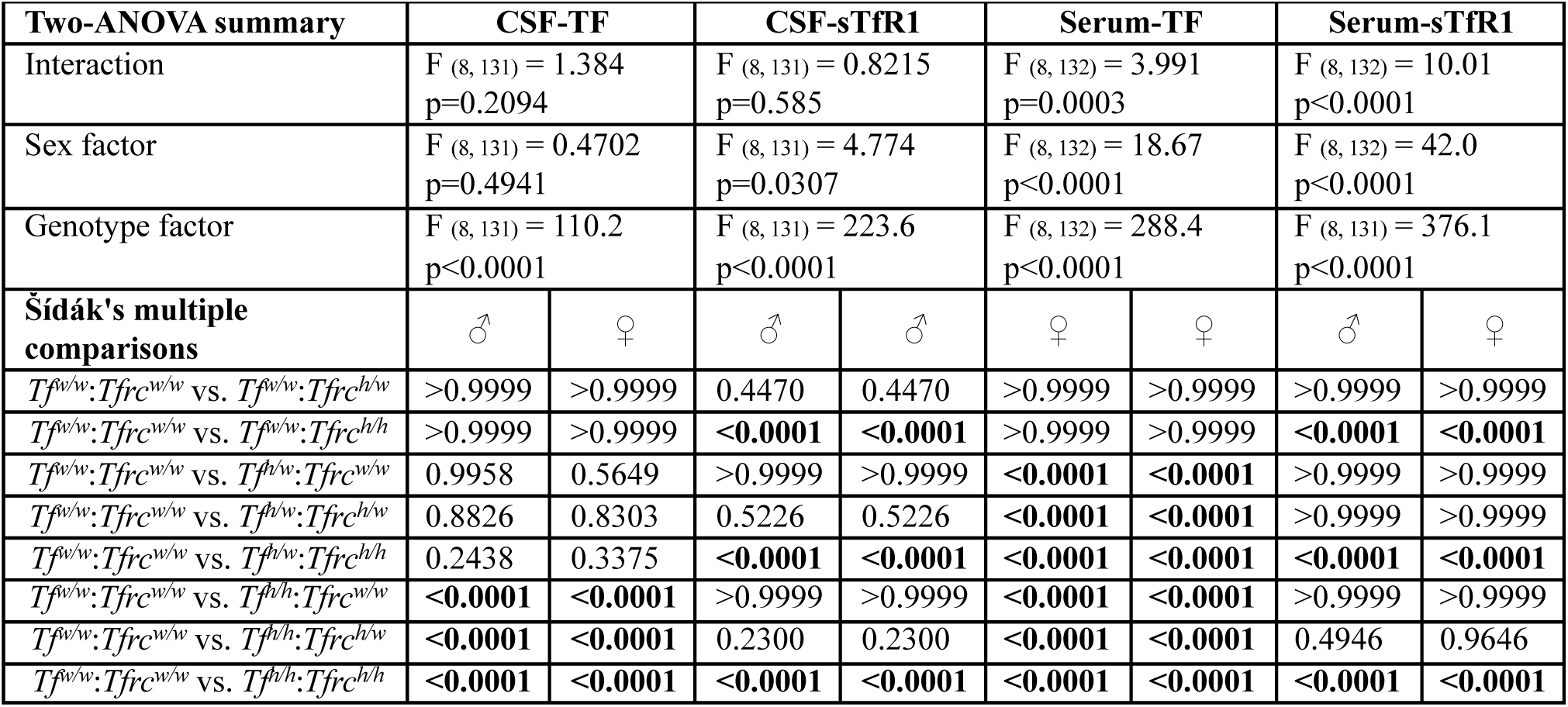
Statistical analysis of CSF and serum TF and sTfR1 levels.

Serum sTfR1 levels in homozygous *Tfrc^h/h^* rats ranged from 2,000 to 3,000 ng/mL (**Figure 4D**). These values fall within or slightly exceed the human reference range of 1,150–2,750 ng/mL (15), suggesting that the humanized TfR1 allele is efficiently expressed, processed, and shed in a physiologically relevant manner. In heterozygous *Tfrc^h/w^*rats, serum levels of human sTfR1 were approximately ninefold lower than those observed in homozygous *Tfrc^h/h^* animals, with a similar pattern observed in CSF (**Figure 4B**). This deviation from the expected gene-dosage effect may reflect impaired formation or reduced cleavage of cross-species TfR1 dimers, from which sTfR1 is generated at the cell surface by metalloproteases.

We were unable to identify well-established reference ranges for Tf and sTfr1 concentrations in wild-type rats, preventing direct comparison to endogenous rat physiology. Nevertheless, these results confirm that the humanized *Tf* and *Tfrc* alleles drive genotype-dependent expression of the human proteins. While human sTfR1 achieves serum levels comparable to those observed in humans, human TF is present at substantially lower concentrations.

### Human TF and TfR1 expression and detection of cross-species TfR1 dimers in KI rats

To further validate expression of the human TF and TfR1 proteins in the KI models, we examined cortical brain lysates, a region highly relevant to the intended application of these models. These experiments aimed to determine whether the humanized alleles drive exclusive expression of the human protein, and whether human TfR1 can form disulfide-linked homodimers as well as cross-species dimers with rat TfR1 in heterozygous animals.

Western blot analysis was performed on cortical lysates from male and female rats of all nine *Tf* and *Tfrc* genotypes **(Figure 5A**). Human TF protein was detected in *Tf^h/w^* and *Tf^h/h^* animals, with the highest expression observed in the homozygous *Tf^h/h^* genotype. Rat Tf was present in *Tf^w/w^* and *Tf^h/w^* animals, but absent in *Tf^h/h^* rats, confirming genotype-dependent expression and reciprocal loss of the endogenous protein in the humanized background.

**Figure 5.**
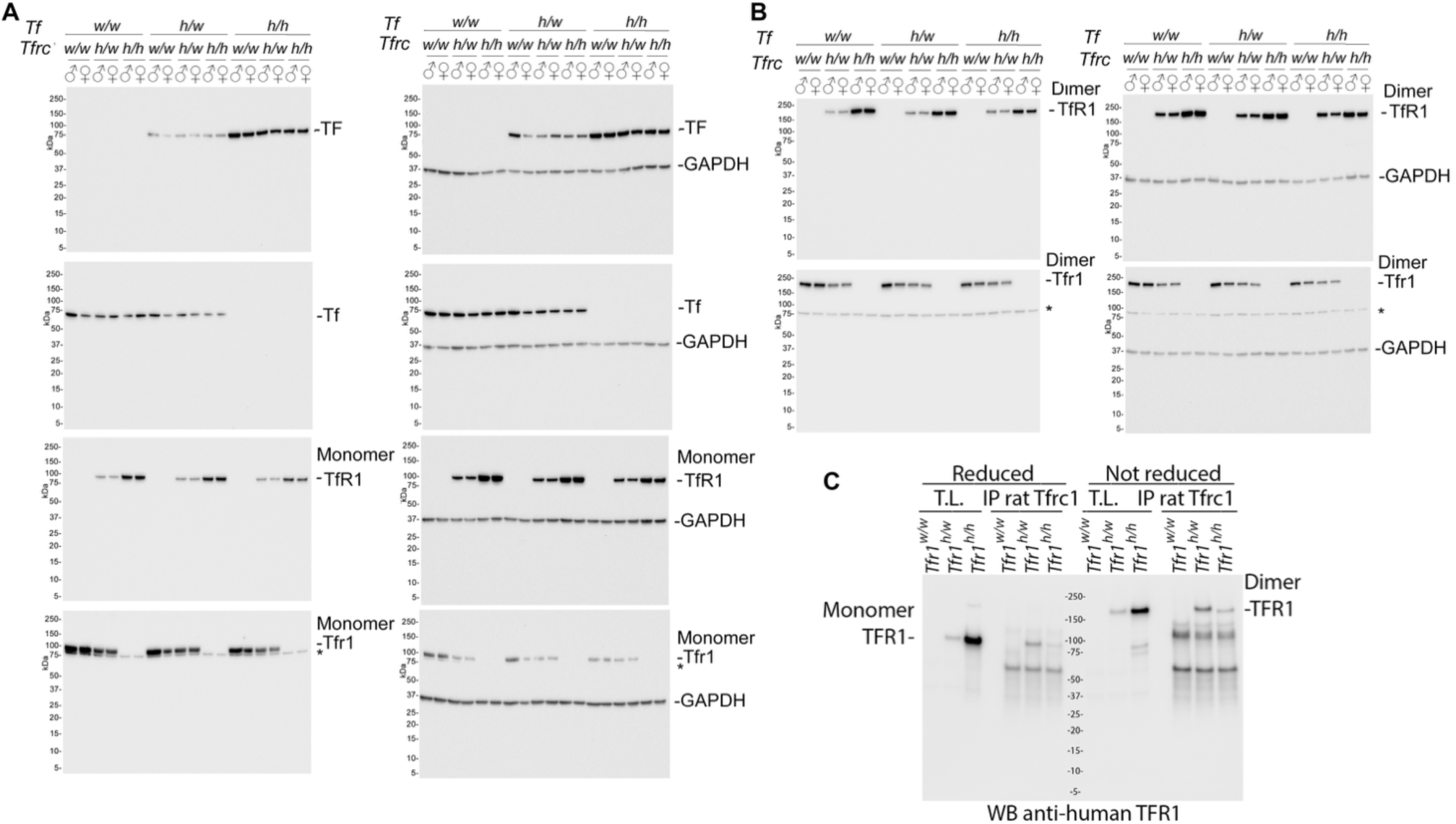
Protein-level validation of human and rat TF/TfR1 expression in brain by Western blot and immunoprecipitation. **A.** From top to bottom, immunoblots show detection of human TF, rat Tf, human TfR1, and rat Tfr1. Human TF and TfR1 were detected only in animals carrying one or two humanized alleles at the respective loci, whereas rat Tf and Tfr1 were present only in animals carrying one or two endogenous (wild-type) alleles. These results confirm species-specific expression of each transgene and absence of antibody cross-reactivity. **B.** Western blot under non-reducing conditions using the same cortical samples as in *A*, prepared without β-mercaptoethanol. A ∼190 kDa band corresponding to dimeric TfR1 was observed. **C.** Immunoprecipitation from cortical lysates using anti-rat Tfr1 antibody followed by Western blot with human-specific TfR1 antibody revealed chimeric dimers composed of human and rat TfR1 in heterozygous *Tfrc^h/w^* rats. All blots were loaded with equal protein amounts. Reducing (with β-mercaptoethanol) and non-reducing conditions were used to assess monomeric and dimeric TfR1 forms, respectively. Asterisks indicate non-specific background bands.

A similar pattern was observed for TfR1. Human TfR1 was detected in *Tfrc^h/w^* and *Tfrc^h/h^* animals, while rat TfR1 was present in *Tfrc^w/w^* and *Tfrc^h/w^* rats but absent in *Tfrc^h/h^* animals, indicating complete replacement by the human allele. In *Tfrc^h/w^* rats, both human and rat TfR1 proteins were co-expressed.

TfR1 functions as a ∼190 kDa homodimer composed of two identical subunits linked by interchain disulfide bonds (9). Under reducing conditions, only the monomeric ∼95 kDa form of human TfR1 was observed (Figure 5A), whereas under non-reducing conditions, a prominent ∼190 kDa band corresponding to disulfide-linked dimers was detected (**Figure 5B**), indicating that the vast majority of human TfR1 exists in a dimeric state. The presence of this dimeric band under non-reducing conditions in homozygous *Tfrc^h/h^* rats suggests efficient formation of disulfide-linked human TfR1 homodimers, likely at the cell surface. In heterozygous *Tfrc^h/w^* animals, these dimers can include both human–human and human–rat TfR1 subunit combinations. While these findings strongly support physiological dimerization of TfR1, we cannot formally exclude the possibility that a fraction of these dimers form post-lysis during sample processing. Additional studies would be required to definitively distinguish in vivo dimerization from potential artifacts introduced during lysis.

To determine whether human and rat TfR1 form cross-species dimers, we performed immunoprecipitations. However, the anti-rat TfR1 antibody used in Figure 5A does not cross-react with human TfR1 in Western blots but showed strong cross-reactivity in immunoprecipitation (IP) assays. Therefore, a different rat-specific TfR1 antibody was used for the IP experiments to increase specificity of the pulldown. This antibody exhibits only partial cross-reactivity with the human protein. This is evidenced by its ability to precipitate small amounts of human TfR1 from *Tfrc^h/h^* brain lysates (**Figure 5C**). However, this antibody pulled down substantially more human TfR1 from *Tfrc^h/w^* rats, despite their lower total expression of the human protein (**Figure 5C**). This disproportionate efficiency strongly suggests that in heterozygous animals, human TfR1 forms disulfide-linked dimers with rat TfR1, allowing the rat-specific antibody to co-precipitate the human protein through its association with the rat TfR1 subunit with high efficiency.

Although we cannot formally exclude the possibility that a fraction of the observed TfR1 dimers formed post-lysis during sample processing, multiple lines of evidence—beyond those shown in Figure 5—support physiological dimerization in vivo. First, the presence of human sTfR1 in the serum of humanized rats, which arises from proteolytic cleavage of mature, membrane-bound TfR1 dimers, indicates proper assembly and surface localization of functional dimeric receptors. Second, humanized TfR1 enables efficient BBB transcytosis of NewroBus constructs (see reference (7) and accompanying paper), further supporting that the human TfR1 is correctly trafficked and functionally active at the BBB.

### Localization of human TF and TfR1 in KI rats

#### Brain

IF staining of cortical sections revealed that anti-human TfR1 antibodies labeled vessel-like structures only in rats carrying at least one humanized *Tfrc* allele, while anti-rat Tfr1 antibodies stained only wild-type *Tfrc* alleles, confirming antibody specificity (**Figure 6A**). In heterozygous *Tfrc^h/w^* rats, co-staining showed that the two proteins co-localized in cortical vasculature, consistent with BBB expression (**Figure 6B**). In contrast, co-staining of human and rat TF proteins in the same slice was not feasible because both anti-human TF and anti-rat Tf antibodies were raised in rabbits. Therefore, adjacent serial sections were stained separately to assess species-specific TF expression. Adjacent serial sections stained for TF showed genotype-dependent expression: rat Tf was present in *Tf^w/w^* and *Tf^h/w^* animals, and human TF was detected only in animals carrying the humanized *Tf* allele, with similar regional distribution, confirming proper brain expression (**Figure 7**).

**Figure 6.**
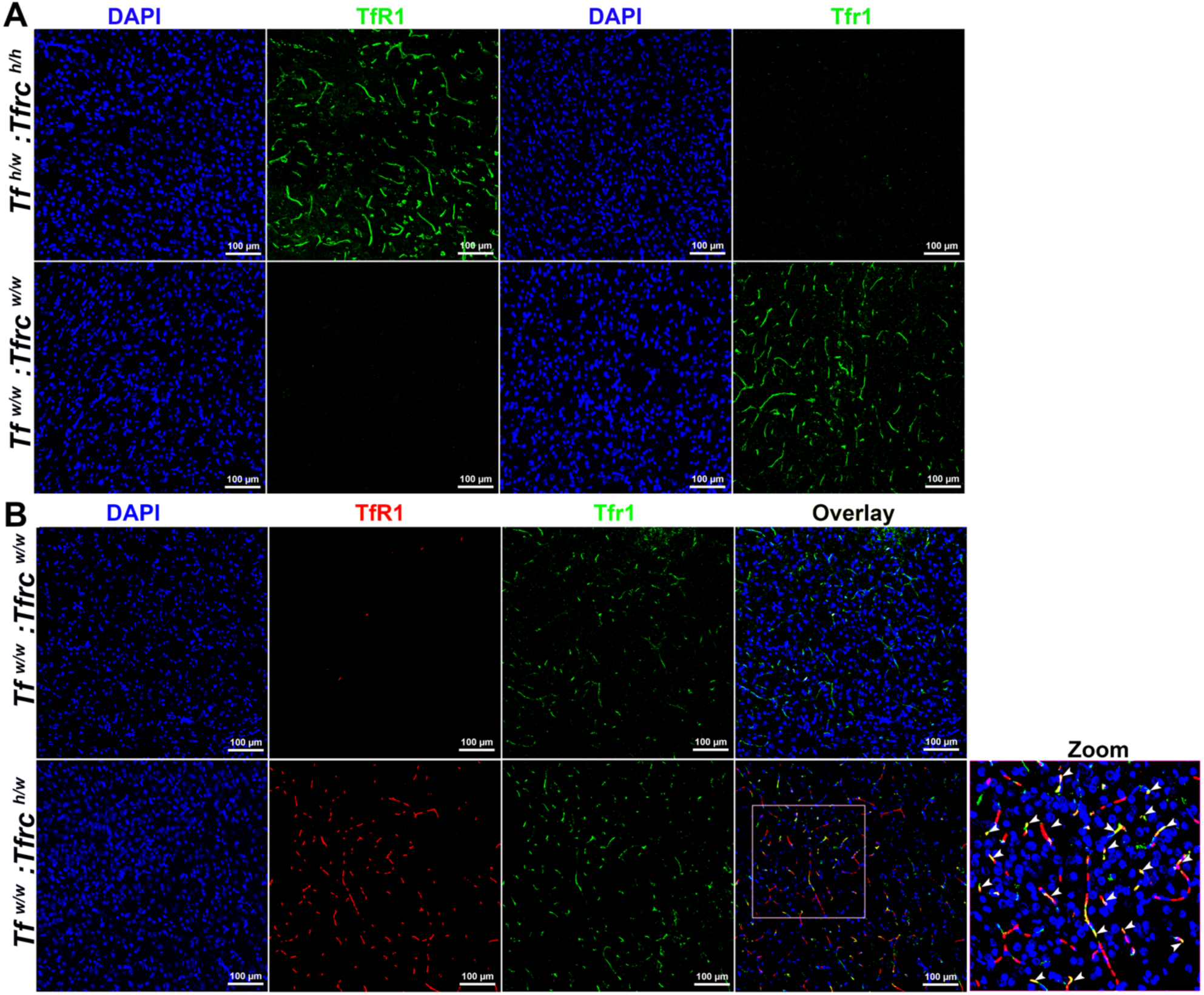
Colocalization of human and rat TfR1 in vessel-like structures of the frontal cortex. **A.** Species-specific immunofluorescence confirms that the anti-human TfR1 antibody (green) detects only human TfR1, while the anti-rat Tfr1 antibody (green) recognizes only the endogenous rat protein. **B.** Dual labeling with anti-human TfR1 (red) and anti-rat Tfr1 (green) in brains of heterozygous *Tfrc^h/w^* rats reveals colocalization in vessel-like structures. No staining was observed with the anti-human TfR1 antibody in *Tfrc^w/w^* rats, confirming its specificity. White arrows in the zoomed-in panel indicate colocalization within the region highlighted by the white square in the overlay.

**Figure 7.**
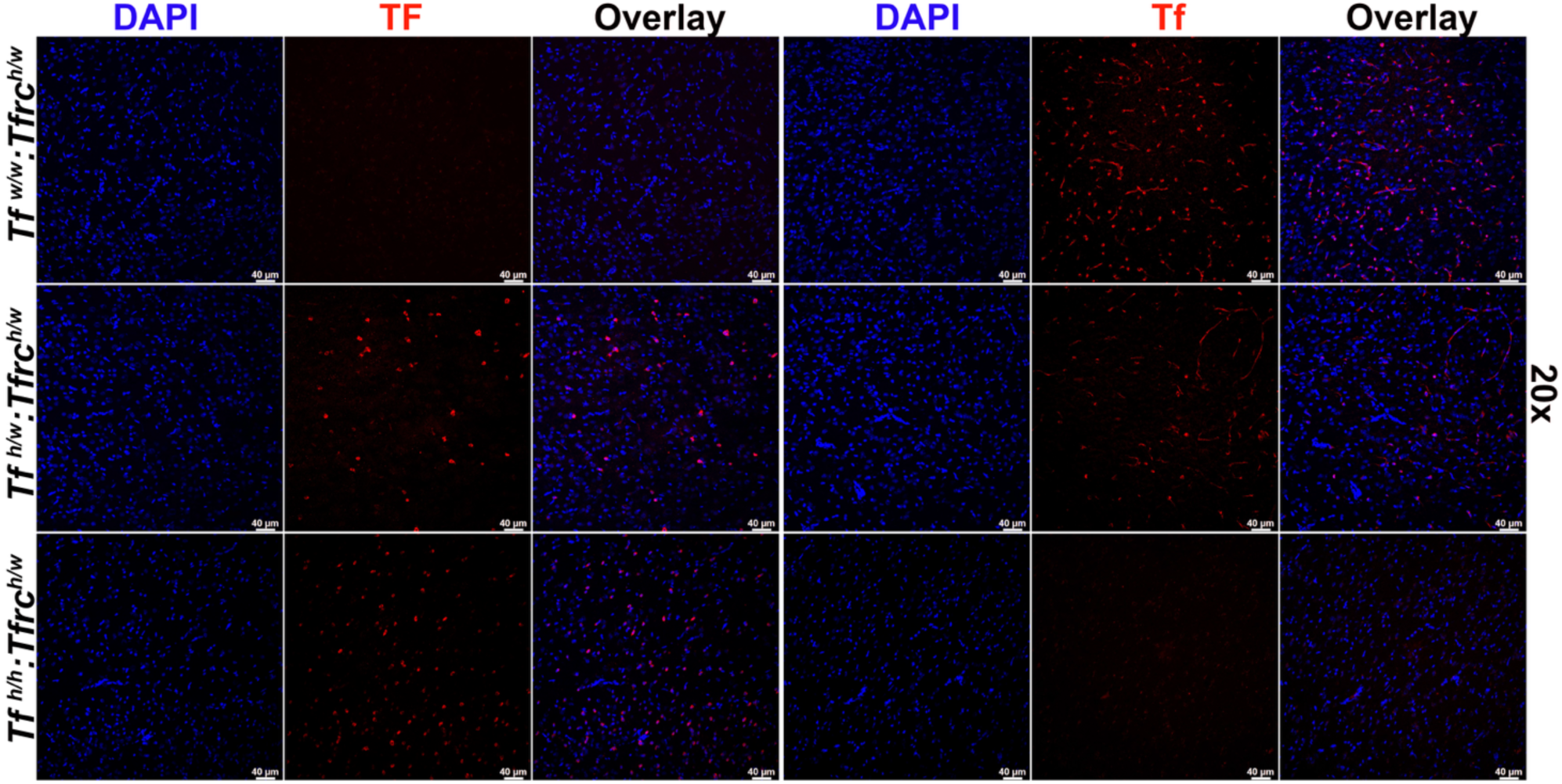
Detection of human and rat transferrin (TF/Tf) in the frontal cortex. Immunofluorescence staining was performed on brain sections from rats with the following genotypes: *Tf^w/w^*, *Tf^h/w^*, and *Tf^h/h^*. Human TF was detected in the frontal cortex of *Tf^h/w^* and *Tf^h/h^* rats, but not in wild-type *Tf^w/w^*animals, confirming human-specific expression. Rat Tf was detected in all genotypes carrying at least one endogenous allele.

#### Duodenum

Rat Tfr1 was localized to villus epithelial cells in *Tfrc^w/w^* and *Tfrc^h/w^* rats, while human Tfr1 was detected in *Tfrc^h/w^* and *Tfrc^h/h^* animals (**Figure 8**). In heterozygous animals, both proteins co-localized along the basolateral membrane, with punctate cytoplasmic staining suggestive of vesicular trafficking. This spatial distribution aligns with known sites of receptor-mediated iron uptake (16). Human TF was observed in villus epithelial cells of *Tf^h/w^* and *Tf^h/h^* rats, consistent with known sites of transferrin synthesis and function. Adjacent sections confirmed complementary expression of rat Tf in *Tf^w/w^* and *Tf^h/w^* animals (**Figure 9**).

**Figure 8.**
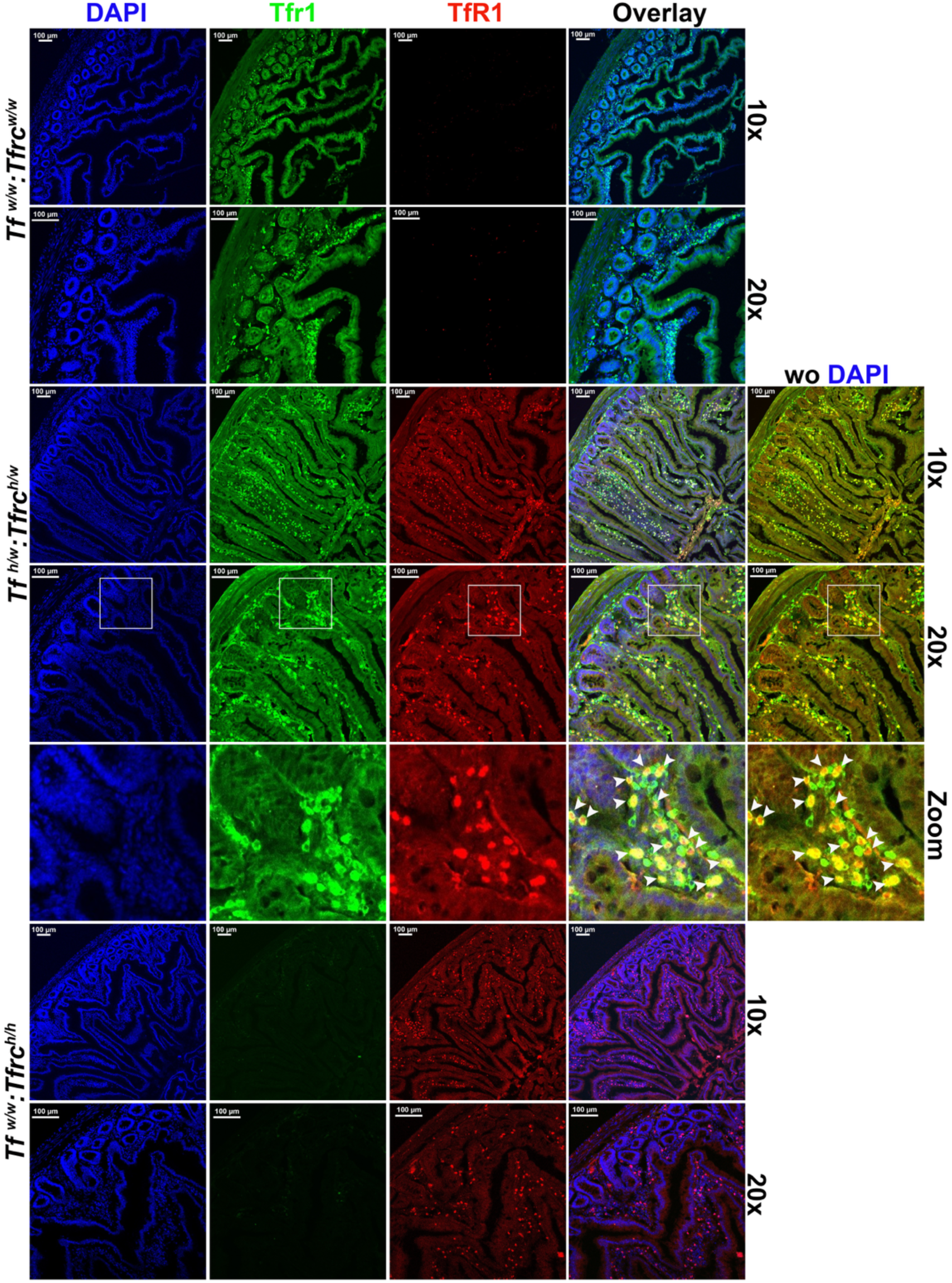
Co-localization of human TfR1 and rat Tfr1 in the duodenum. Sections from *Tfrc^w/w^*, *Tfrc^h/w^*, and *Tfrc^h/h^* rats were co-stained with antibodies specific for human TfR1 and rat Tfr1. Co-localization was observed only in heterozygous *Tfrc^h/w^* rats, while *Tfrc^w/w^* and *Tfrc^h/h^* rats showed exclusive expression of rat or human TfR1, respectively. White arrows in the zoomed-in panel indicate co-localization spots within the region highlighted by the white square in the overlay panel.

**Figure 9.**
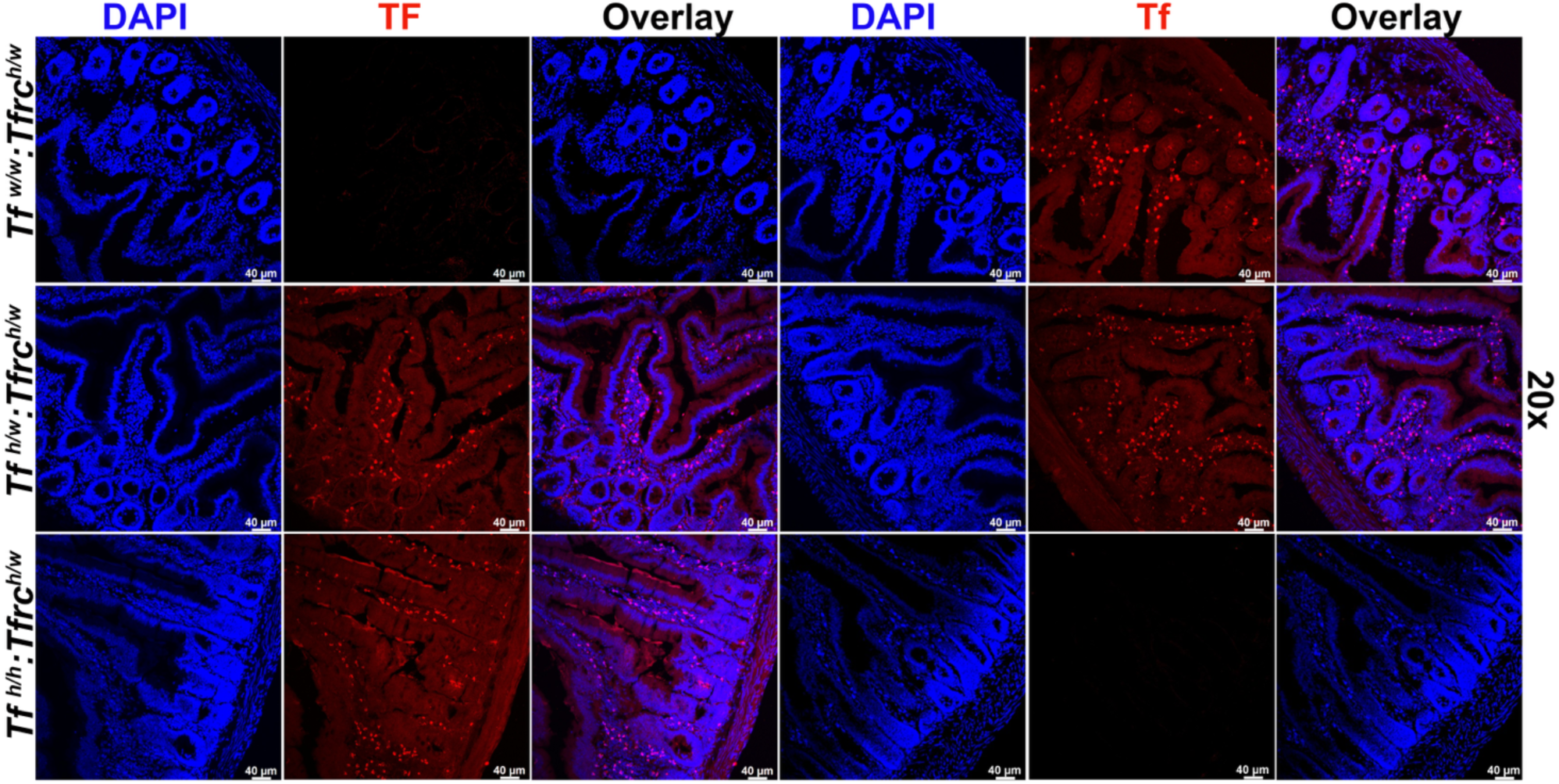
Detection of human and rat transferrin (TF/Tf) in the duodenum. Immunostaining revealed human TF expression in *Tf^h/w^*and *Tf^h/h^* genotypes, with no signal in *Tf^w/w^* animals. Rat Tf was detected in all genotypes carrying at least one endogenous allele.

#### Kidney

In the kidney, rat Tfr1 was detected in tubular epithelial cells of *Tfrc^w/w^* and *Tfrc^h/w^* rats, while human TfR1 was present in *Tfrc^h/w^* and *Tfrc^h/h^* rats (**Figure 10**). Co-localization was seen in heterozygous animals, especially along the basolateral membrane of tubules. This spatial distribution is consistent with known TfR1 localization in proximal tubules, where receptor-mediated iron uptake occurs (17). Rat Tf was expressed in *Tf^w/w^* and *Tf^h/w^* rats, while human TF was seen in *Tf^h/w^* and *Tf^h/h^* animals. TF staining was cytoplasmic and confined to tubular cells, with no signal in glomeruli (**Figure 11**).

**Figure 10.**
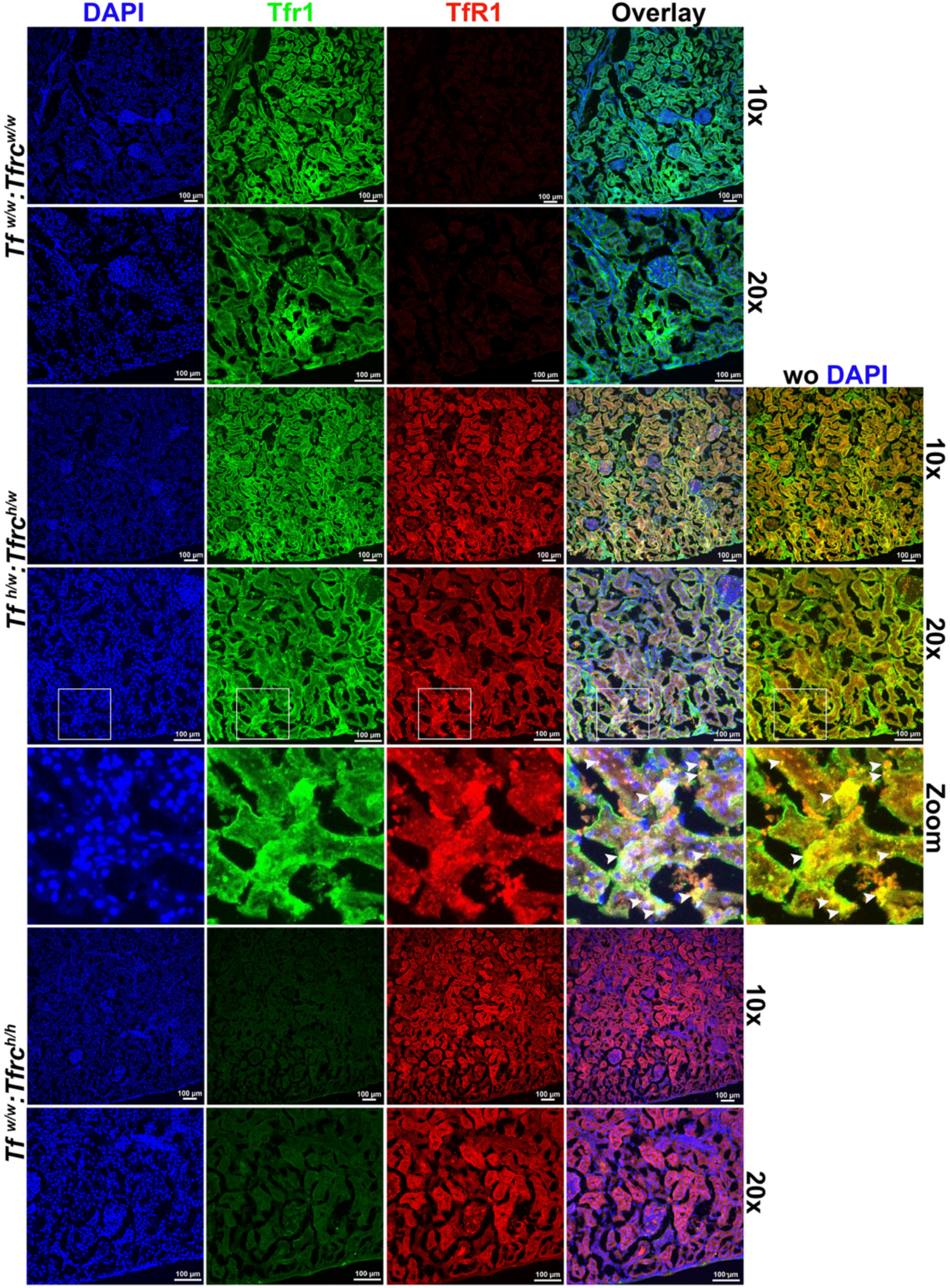
Co-localization of human TfR1 and rat Tfr1 in the kidney. Kidney sections from *Tfrc^w/w^*, *Tfrc^h/w^*, and *Tfrc^h/h^*rats were stained for both human TfR1 and rat Tfr1. Co-localization was observed only in *Tfrc^h/w^* rats, while *Tfrc^w/w^* and *Tfrc^h/h^*showed signal only for rat or human TfR1, respectively. White arrows in the zoomed-in panel indicate co-localization spots within the region highlighted by the white square in the overlay panel.

**Figure 11.**
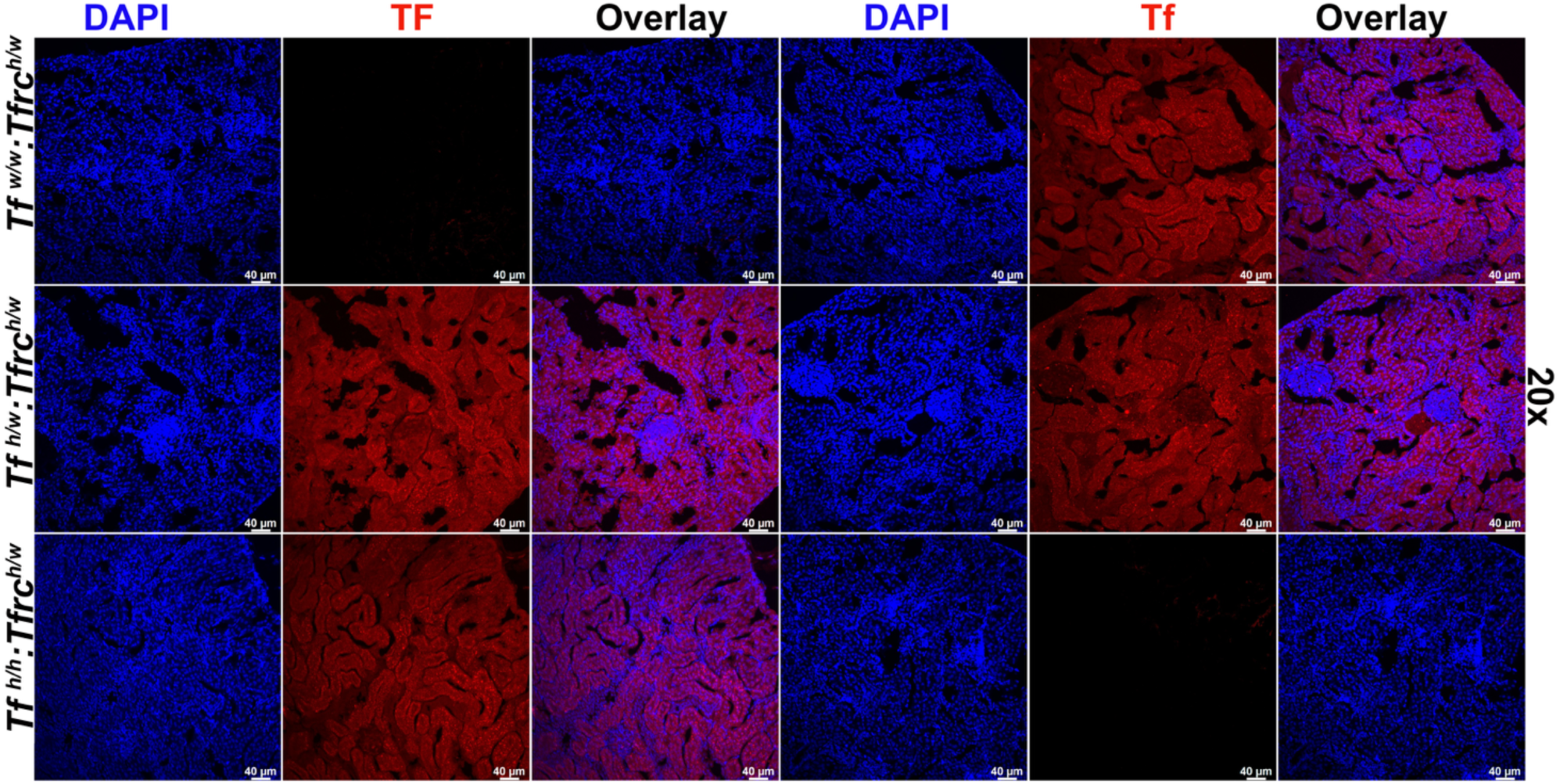
Detection of human and rat transferrin (TF/Tf) in the kidney. Human TF was detected in *Tf^h/w^* and *Tf^h/h^* genotypes, but not in *Tf^w/w^* animals. Rat Tf was detected in all animals carrying wild-type *Tf* alleles.

#### Liver

Rat Tfr1 was found in periportal hepatocytes of *Tfrc^w/w^*and *Tfrc^h/w^* animals, and human TfR1 was similarly localized in *Tfrc^h/w^* and *Tfrc^h/h^* livers (**Figure 12**). In heterozygotes, co-localization of both ortholog was observed along the plasma membrane of periportal hepatocytes, where iron uptake is highest (18). Rat Tf was detected in *Tf^w/w^* and *Tf^h/w^* animals; human TF was expressed in hepatocytes of *Tf^h/w^* and *Tf^h/h^* rats, with cytoplasmic, periportal staining consistent with synthesis and secretion (**Figure 13**).

**Figure 12.**
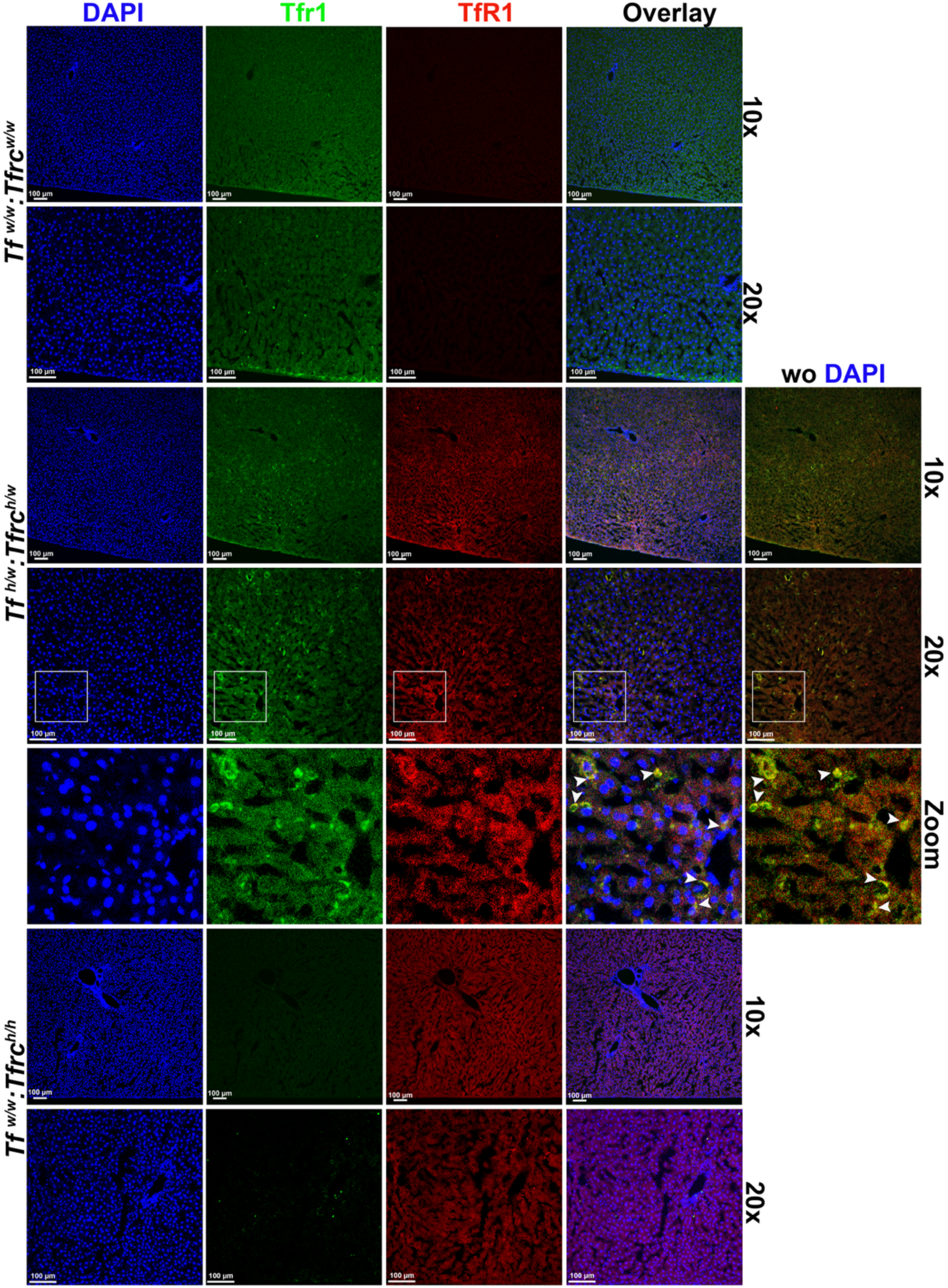
Co-localization of human TfR1 and rat Tfr1 in the liver. Liver sections from *Tfrc^w/w^*, *Tfrc^h/w^*, and *Tfrc^h/h^*genotypes were stained with species-specific antibodies. Co-localization of human TfR1 and rat Tfr1 was observed only in *Tfrc^w/w^* rats. White arrows in the zoomed-in panel indicate regions of co-localization.

**Figure 13.**
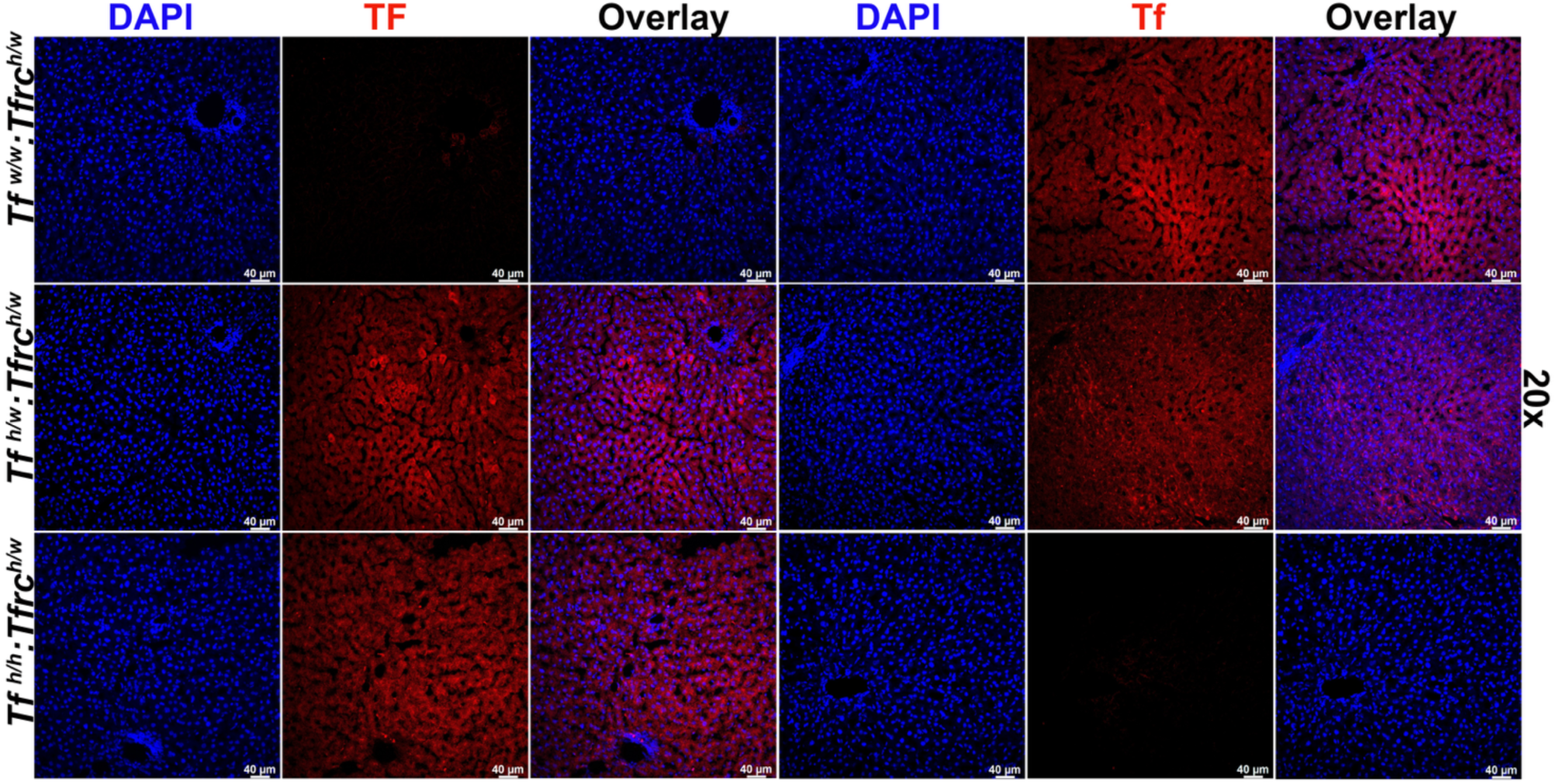
Detection of human and rat transferrin (TF/Tf) in the liver. Human TF was present in *Tf^h/w^* and *Tf^h/h^* rats, but absent in *Tf^w/w^* animals. Rat Tf was detected in animals with wild-type *Tf* alleles.

#### Lung

Rat TfR1 was localized to alveolar and airway epithelium in *Tfrc^w/w^*and *Tfrc^h/w^* rats, while human TfR1 was detected in the same structures in *Tfrc^h/w^* and *Tfrc^h/h^* animals (**Figure 14**). Co-localization in heterozygous rats supported appropriate spatial distribution. TF staining showed alveolar and airway localization, with rat and human Tf expressed according to genotype, and no major differences in intensity between heterozygous and homozygous humanized rats (**Figure 15**). The presence of both proteins in alveolar parenchyma and airway epithelium is consistent with the role of TfR1 in supporting iron uptake for respiratory and immune cell function in pulmonary tissues (19).

**Figure 14.**
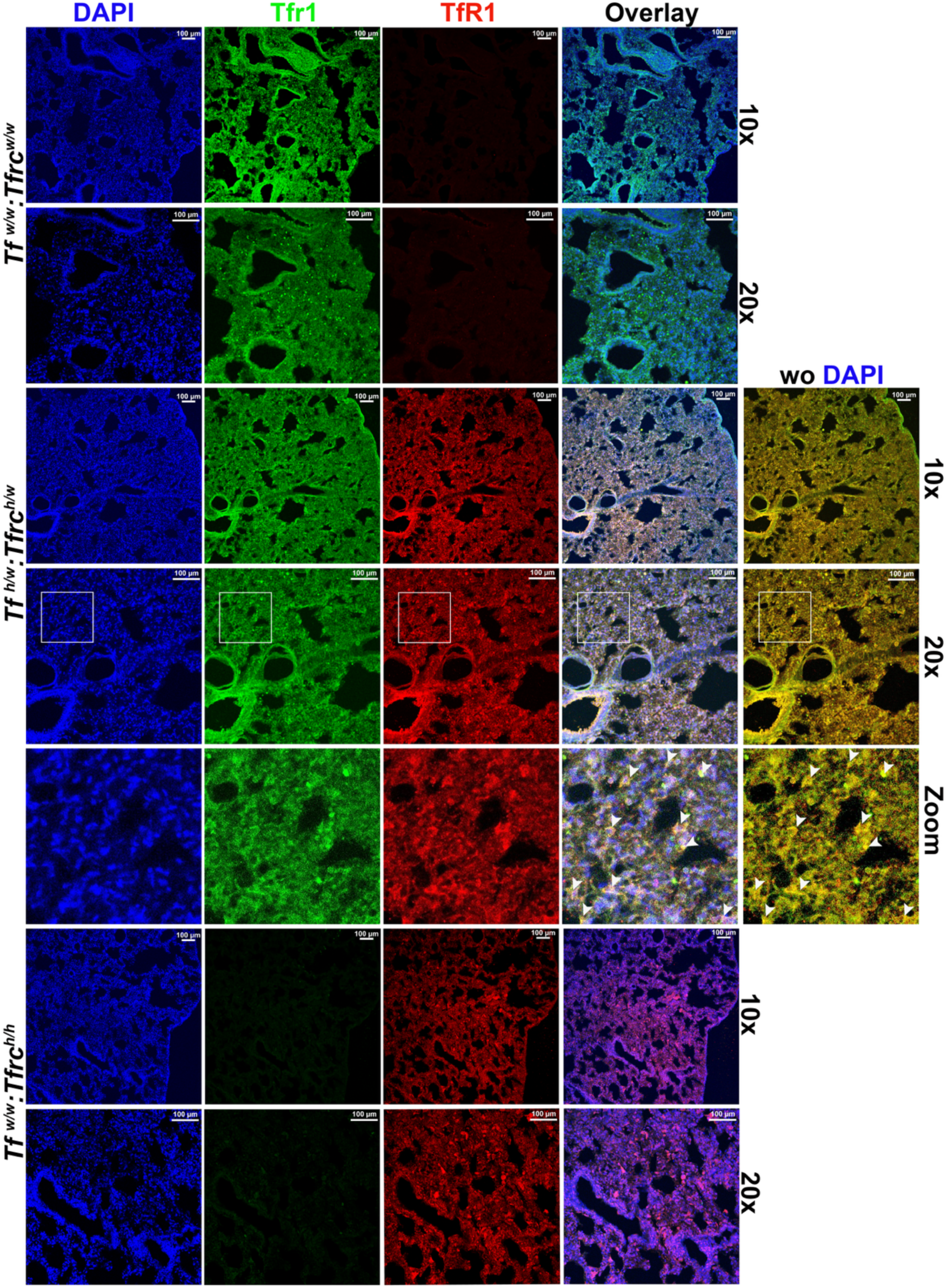
Co-localization of human TfR1 and rat Tfr1 in the lung. Lung sections from *Tfrc^w/w^*, *Tfrc^h/w^*, and *Tfrc^h/h^* rats were stained with species-specific antibodies. Co-localization was observed only in *Tfrc^w/w^* rats. White arrows highlight the co-localization sites within the region demarcated by the white square.

**Figure 15.**
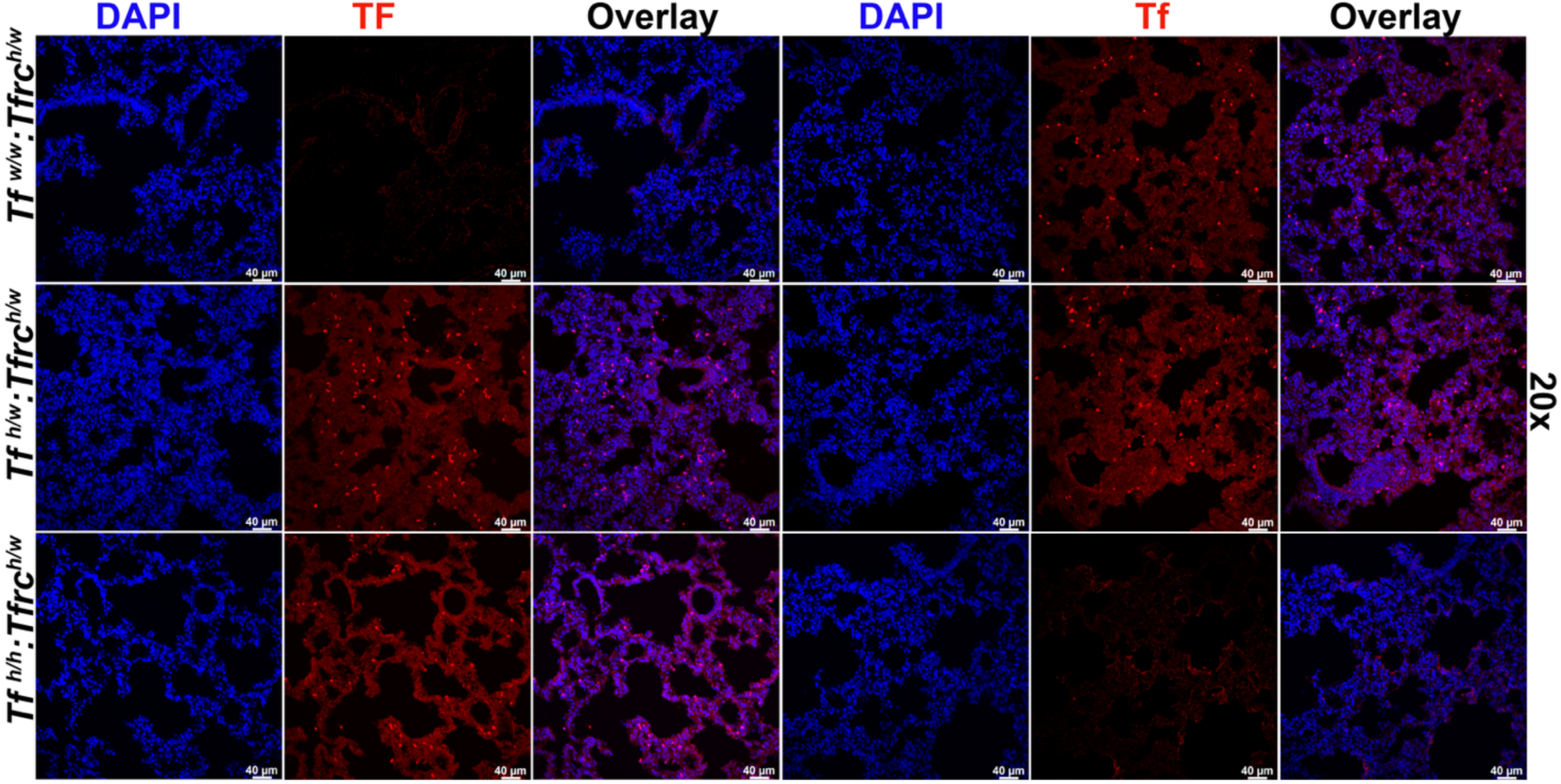
Detection of human and rat transferrin (TF/Tf) in the lung. Human TF expression was detected in *Tf^h/w^* and *Tf^h/h^* rats, but not in *Tf^w/w^* animals. Rat Tf staining was present in genotypes retaining at least one wild-type allele.

In summary, the humanized *Tf* and *Tfrc* alleles are correctly expressed and human TF and TfR1 proteins localized in all major iron-regulatory tissues, including brain, duodenum, kidney, liver, and lung. Their distribution mirrors that of the endogenous rat proteins, with co-localization of human and rat TfR1 supporting physiological expression and potential cross-species dimerization.

### Complete blood count (CBC) analysis

To assess whether the humanized *Tf* and *Tfrc* alleles support normal systemic iron metabolism and hematopoiesis, we performed CBC analysis on 8-week-old male and female rats representing all genotypes: *Tf^w/w^:Tfrc^w/w^*, *Tf^w/w^:Tfrc^h/w^*, *Tf^w/w^:Tfrc^h/h^*, *Tf^h/w^:Tfrc^w/w^*, *Tf^h/w^:Tfrc^h/w^*, *Tf^h/w^:Tfrc^h/h^*, *Tf^h/h^:Tfrc^w/w^*, *Tf^h/h^:Tfrc^h/w^*, *Tf^h/h^:Tfrc^h/h^*(**Figure 16**). Blood samples were analyzed using an automated hematology analyzer. The CBC analysis included standard leukocyte, erythrocyte, and platelet parameters. These included total and differential white blood cell count (neutrophils, lymphocytes, monocytes, eosinophils, and basophils, both absolute and percentage), red blood cell indices (RBC count, hemoglobin, hematocrit, MCV, MCH, MCHC, and RDW), and platelet measures (platelet count and mean platelet volume). To maintain clarity and avoid overcomplicating graphical representations, the full statistical analysis is provided in Tables 2-4.

**Figure 16.**
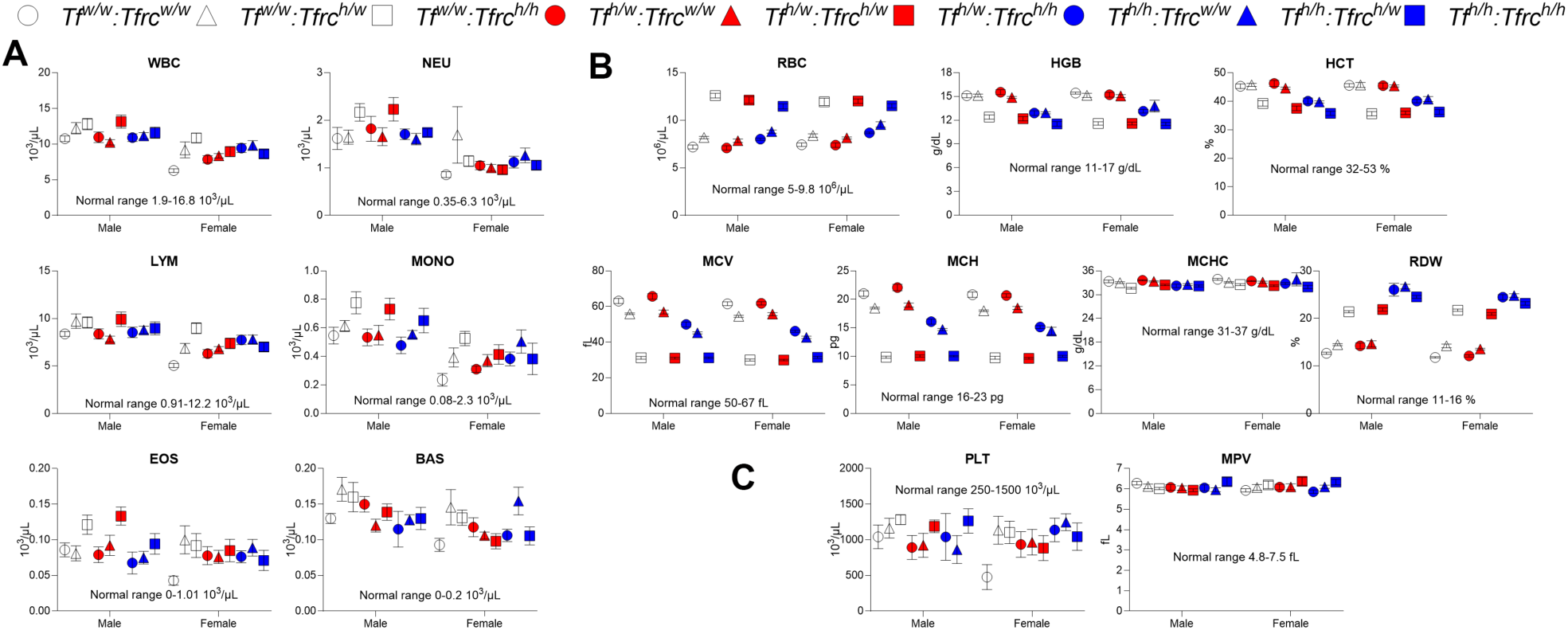
CBC analysis across all nine genotypic combinations of humanized *Tf* and *Tfrc* knock-in rats. **A.** Leukocyte parameters—white blood cells (WBC), neutrophils (NEU), lymphocytes (LYM), monocytes (MONO), eosinophils (EOS), and basophils (BAS)—are shown by genotype and sex, with normal physiological ranges indicated for reference. Two-way ANOVA revealed significant effects of genotype and sex, with a significant genotype × sex interaction specifically for NEU. Post hoc comparisons are reported in Table 2. **B.** Erythrocyte parameters—including red blood cell count (RBC), hemoglobin (HGB), hematocrit (HCT), mean corpuscular volume (MCV), mean corpuscular hemoglobin (MCH), mean corpuscular hemoglobin concentration (MCHC), and red cell distribution width (RDW)—are plotted across genotypes and sexes. Significant genotype effects were detected for all parameters (see Table 3 for full statistical details). **C.** Platelet parameters—platelet count (PLT) and mean platelet volume (MPV)—were unaffected by genotype, sex, or their interaction, as determined by two-way ANOVA (see Table 4). All data are presented as mean ± SEM. Normal reference ranges are indicated by shaded areas in each plot, where applicable.

**Table 2.**
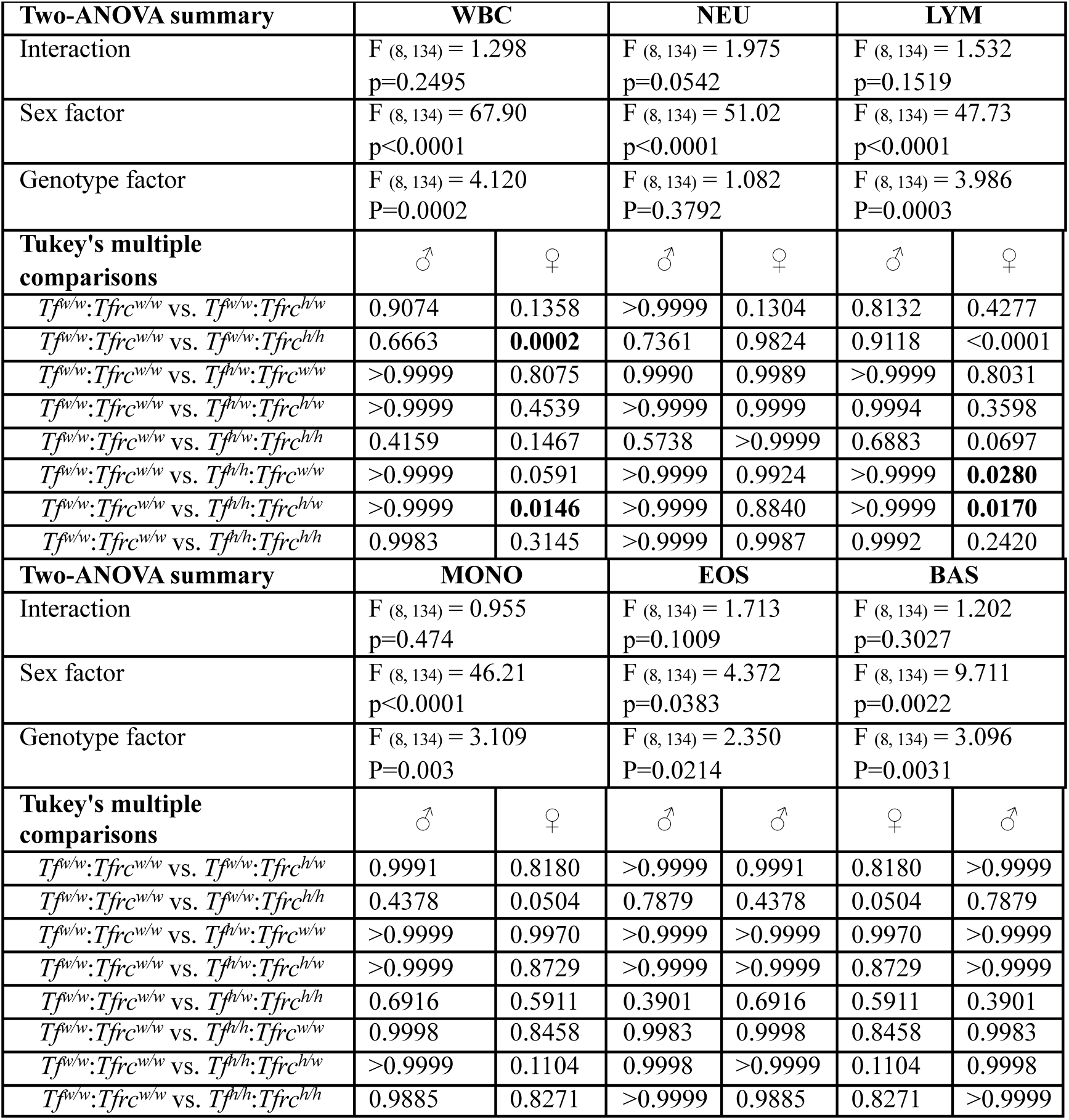
Statistical analysis of WBC shown in Figure 16A.

**Table 3.**
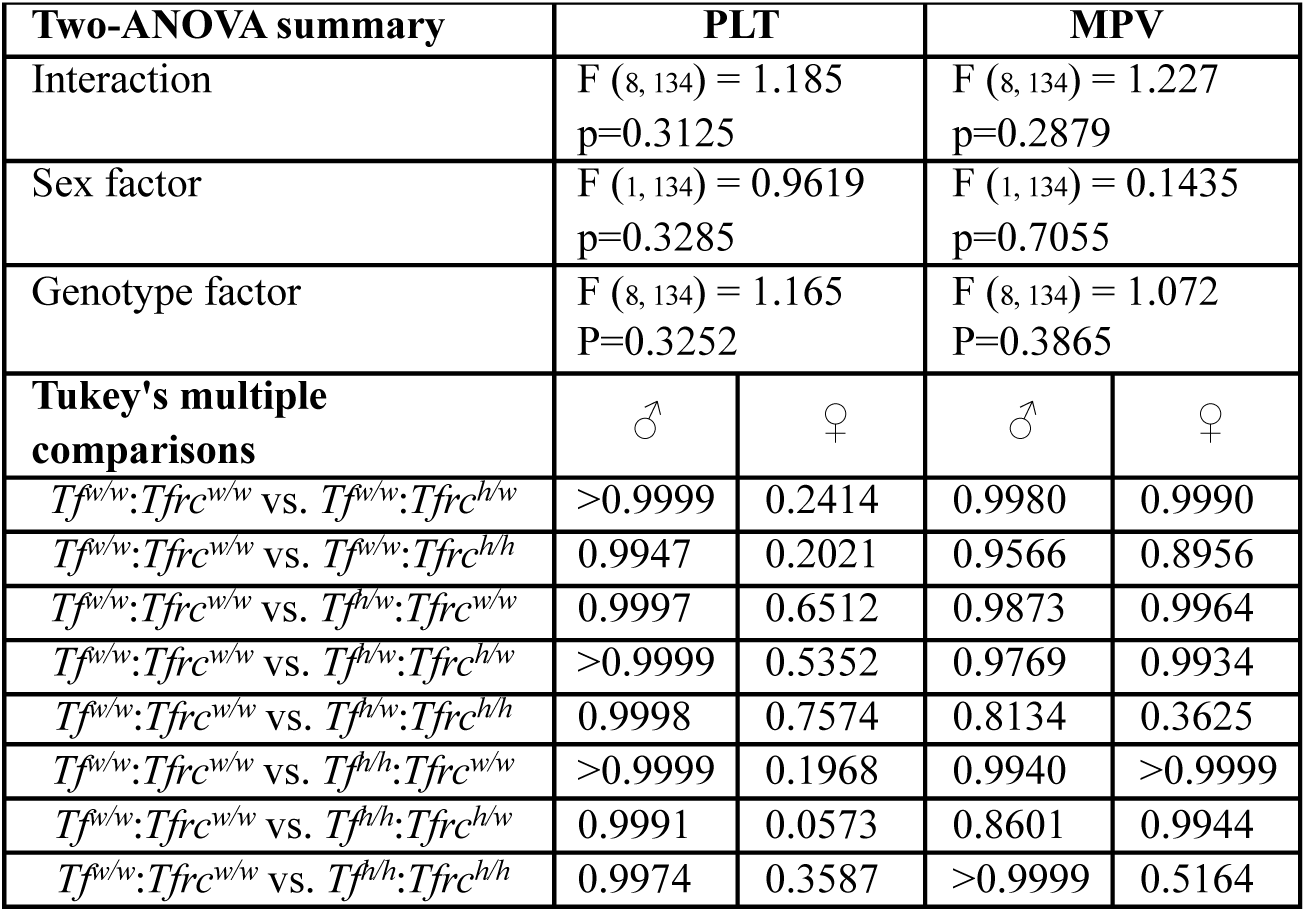
Statistical analysis of PLT shown in Figure 16B.

**Table 4.**
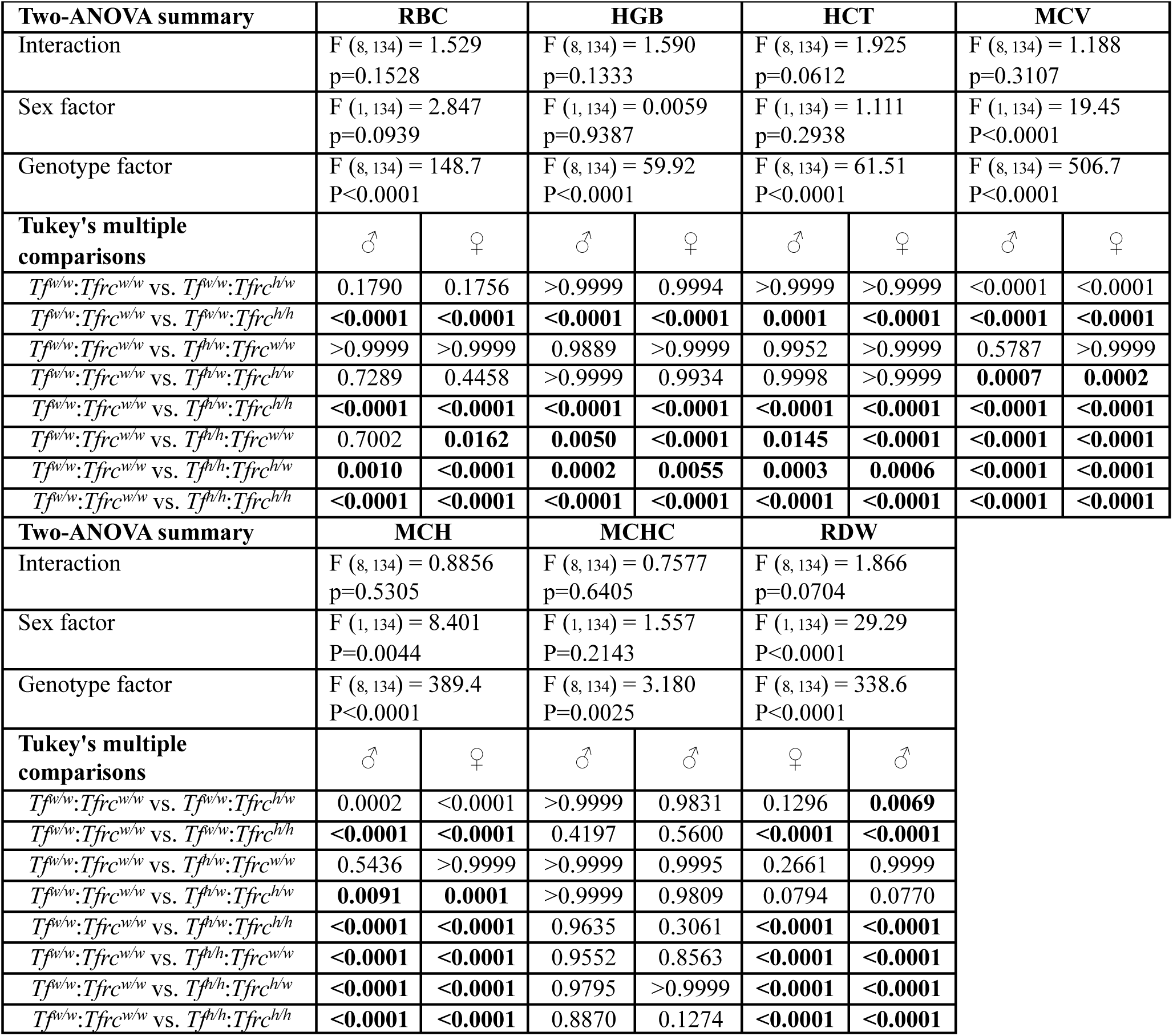
Statistical analysis of RBC shown in Figure 16C.

| **Two-ANOVA summary** | **RBC** |  | **HGB** |  | **HCT** |  | **MCV** |  |
| --- | --- | --- | --- | --- | --- | --- | --- | --- |
| Interaction | F (8, 134) = 1.529<br>p=0.1528 |  | F (8, 134) = 1.590<br>p=0.1333 |  | F (8, 134) = 1.925<br>p=0.0612 |  | F (8, 134) = 1.188<br>p=0.3107 |  |
| Sex factor | F (1, 134) = 2.847<br>p=0.0939 |  | F (1, 134) = 0.0059<br>p=0.9387 |  | F (1, 134) = 1.111<br>p=0.2938 |  | F (1, 134) = 19.45<br>P<0.0001 |  |
| Genotype factor | F (8, 134) = 148.7<br>P<0.0001 |  | F (8, 134) = 59.92<br>P<0.0001 |  | F (8, 134) = 61.51<br>P<0.0001 |  | F (8, 134) = 506.7<br>P<0.0001 |  |
| **Tukey's multiple comparisons** | ♂ | ♀ | ♂ | ♀ | ♂ | ♀ | ♂ | ♀ |
| $Tf^{w/w}:Tfrc^{w/w}$ vs. $Tf^{w/w}:Tfrc^{h/w}$ | 0.1790 | 0.1756 | >0.9999 | 0.9994 | >0.9999 | >0.9999 | <0.0001 | <0.0001 |
| $Tf^{w/w}:Tfrc^{w/w}$ vs. $Tf^{w/w}:Tfrc^{h/h}$ | <0.0001 | <0.0001 | <0.0001 | <0.0001 | 0.0001 | <0.0001 | <0.0001 | <0.0001 |
| $Tf^{w/w}:Tfrc^{w/w}$ vs. $Tf^{h/w}:Tfrc^{w/w}$ | >0.9999 | >0.9999 | 0.9889 | >0.9999 | 0.9952 | >0.9999 | 0.5787 | >0.9999 |
| $Tf^{w/w}:Tfrc^{w/w}$ vs. $Tf^{h/w}:Tfrc^{h/w}$ | 0.7289 | 0.4458 | >0.9999 | 0.9934 | 0.9998 | >0.9999 | 0.0007 | 0.0002 |
| $Tf^{w/w}:Tfrc^{w/w}$ vs. $Tf^{h/w}:Tfrc^{h/h}$ | <0.0001 | <0.0001 | <0.0001 | <0.0001 | <0.0001 | <0.0001 | <0.0001 | <0.0001 |
| $Tf^{w/w}:Tfrc^{w/w}$ vs. $Tf^{h/h}:Tfrc^{w/w}$ | 0.7002 | 0.0162 | 0.0050 | <0.0001 | 0.0145 | <0.0001 | <0.0001 | <0.0001 |
| $Tf^{w/w}:Tfrc^{w/w}$ vs. $Tf^{h/h}:Tfrc^{h/w}$ | 0.0010 | <0.0001 | 0.0002 | 0.0055 | 0.0003 | 0.0006 | <0.0001 | <0.0001 |
| $Tf^{w/w}:Tfrc^{w/w}$ vs. $Tf^{h/h}:Tfrc^{h/h}$ | <0.0001 | <0.0001 | <0.0001 | <0.0001 | <0.0001 | <0.0001 | <0.0001 | <0.0001 |
| **Two-ANOVA summary** | **MCH** |  | **MCHC** |  | **RDW** |  |  |  |
| Interaction | F (8, 134) = 0.8856<br>p=0.5305 |  | F (8, 134) = 0.7577<br>p=0.6405 |  | F (8, 134) = 1.866<br>p=0.0704 |  |  |  |
| Sex factor | F (1, 134) = 8.401<br>P=0.0044 |  | F (1, 134) = 1.557<br>P=0.2143 |  | F (1, 134) = 29.29<br>P<0.0001 |  |  |  |
| Genotype factor | F (8, 134) = 389.4<br>P<0.0001 |  | F (8, 134) = 3.180<br>P=0.0025 |  | F (8, 134) = 338.6<br>P<0.0001 |  |  |  |
| **Tukey's multiple comparisons** | ♂ | ♀ | ♂ | ♂ | ♀ | ♂ |  |  |
| $Tf^{w/w}:Tfrc^{w/w}$ vs. $Tf^{w/w}:Tfrc^{h/w}$ | 0.0002 | <0.0001 | >0.9999 | 0.9831 | 0.1296 | 0.0069 | | |
| $Tf^{w/w}:Tfrc^{w/w}$ vs. $Tf^{w/w}:Tfrc^{h/h}$ | <0.0001 | <0.0001 | 0.4197 | 0.5600 | <0.0001 | <0.0001 | | |
| $Tf^{w/w}:Tfrc^{w/w}$ vs. $Tf^{h/w}:Tfrc^{w/w}$ | 0.5436 | >0.9999 | >0.9999 | 0.9995 | 0.2661 | 0.9999 | | |
| $Tf^{w/w}:Tfrc^{w/w}$ vs. $Tf^{h/w}:Tfrc^{h/w}$ | 0.0091 | 0.0001 | >0.9999 | 0.9809 | 0.0794 | 0.0770 | | |
| $Tf^{w/w}:Tfrc^{w/w}$ vs. $Tf^{h/w}:Tfrc^{h/h}$ | <0.0001 | <0.0001 | 0.9635 | 0.3061 | <0.0001 | <0.0001 | | |
| $Tf^{w/w}:Tfrc^{w/w}$ vs. $Tf^{h/h}:Tfrc^{w/w}$ | <0.0001 | <0.0001 | 0.9552 | 0.8563 | <0.0001 | <0.0001 | | |
| $Tf^{w/w}:Tfrc^{w/w}$ vs. $Tf^{h/h}:Tfrc^{h/w}$ | <0.0001 | <0.0001 | 0.9795 | >0.9999 | <0.0001 | <0.0001 | | |
| $Tf^{w/w}:Tfrc^{w/w}$ vs. $Tf^{h/h}:Tfrc^{h/h}$ | <0.0001 | <0.0001 | 0.8870 | 0.1274 | <0.0001 | <0.0001 | | |

Two-way ANOVA revealed a strong sex effect across all leukocyte parameters, with significant differences between males and females in total WBCs, neutrophils, lymphocytes, monocytes, eosinophils, and basophils (**Figure 16A**, statistical comparison is in Table 2). In contrast, genotype effects were modest and limited to specific cell types, with no significant differences observed in neutrophil counts (p = 0.38) and only mild effects on other leukocyte subsets (e.g., lymphocytes p = 0.0003, monocytes p = 0.003, basophils p = 0.0031). Significant genotype-by-sex interactions were not detected for any parameter (all p > 0.05). Post hoc Tukey’s tests confirmed that most genotype comparisons to wild-type controls were non-significant, with a few isolated differences in females. Overall, leukocyte profiles were primarily shaped by sex, and genotype-dependent alterations were few and subtle, suggesting that peripheral immune cell composition is largely preserved across all humanized genotypes.

Similarly, there were no significant differences in any platelet parameters (PLT, MPV) based on genotype or sex. This suggests that platelet production and morphology remain unaffected in the *Tf* and *Tfrc* knock-in models under baseline conditions (**Figure 16B**, statistical comparison is in Table 3).

Strong genotype-dependent effects were instead observed in red blood cell indices. Two-way ANOVA showed significant genotype effects on all erythrocyte parameters, while sex was a significant factor for MCV, MCH, and RDW (**Figure 16C**, statistical comparison is in Table 4). Rats homozygous for the humanized *Tfrc* allele showed increased RBC counts alongside reductions in HGB, HCT, MCV, and MCH, as well as elevated RDW, relative to *Tfrc*^w/w^ controls. These effects were consistent across sexes and independent of *Tf* genotype. Similarly, *Tf*^h/h^ rats displayed lower HGB, HCT, MCV, MCH, and MCHC, and higher RDW compared to wild-type controls, indicating that humanization of either *Tf* or *Tfrc* in the homozygous state alters red blood cell morphology. The observed pattern—microcytosis, hypochromia, and increased size variability—suggests ineffective erythropoiesis or impaired iron utilization, potentially due to disrupted iron trafficking.

In contrast, no red blood cell alterations were observed in heterozygous *Tf^h/w^:Tfrc^w/w^* rats, while double heterozygous *Tf^w/w^:Tfrc^h/w^* rats exhibited only mild hematologic changes. RBC, HGB, HCT, and RDW remained within normal limits, suggesting preserved erythropoiesis and iron homeostasis. However, these rats showed reductions in MCV and MCH across sexes, consistent with mild microcytosis and hypochromia. RDW was selectively increased in female heterozygotes *Tf^w/w^:Tfrc^h/w^* suggesting a sex-specific effect on erythrocyte size variability. Double heterozygous *Tf^h/w^:Tfrc^h/w^* rats displayed a similar but attenuated phenotype relative to *Tf^w/w^:Tfrc^h/w^*single heterozygotes. MCV and MCH remained mildly reduced, but the female-specific RDW increase observed in *Tf^w/w^:Tfrc^h/w^*rats was absent, suggesting that the presence of one human *Tf* allele may partially mitigate the effects of *Tfrc* heterozygosity on erythrocyte morphology.

Altogether, these findings demonstrate that while homozygosity for humanized *Tf* or *Tfrc* leads to measurable disruptions in erythropoiesis and iron metabolism, heterozygosity for either or both alleles maintains normal hematopoietic function with only mild, subclinical alterations.

### Tissue iron levels

To determine whether humanization of *Tf* and *Tfrc* alters systemic or CNS iron distribution, we measured non-heme iron levels in dried tissue samples from spleen, liver, serum, heart, and hippocampus in male and female rats across all genotypes. We also calculated the spleen-to-liver iron ratio as a measure of peripheral iron distribution. Iron levels were expressed as µg/g dry tissue and analyzed separately by sex due to significant sex effects and sex × genotype interactions. Two-way ANOVA revealed significant effects of genotype, sex, and their interaction on spleen, liver, spleen/liver ratio, and hippocampus iron levels (**Figure 17**, statistical comparison is in Table 5).

**Figure 17.**
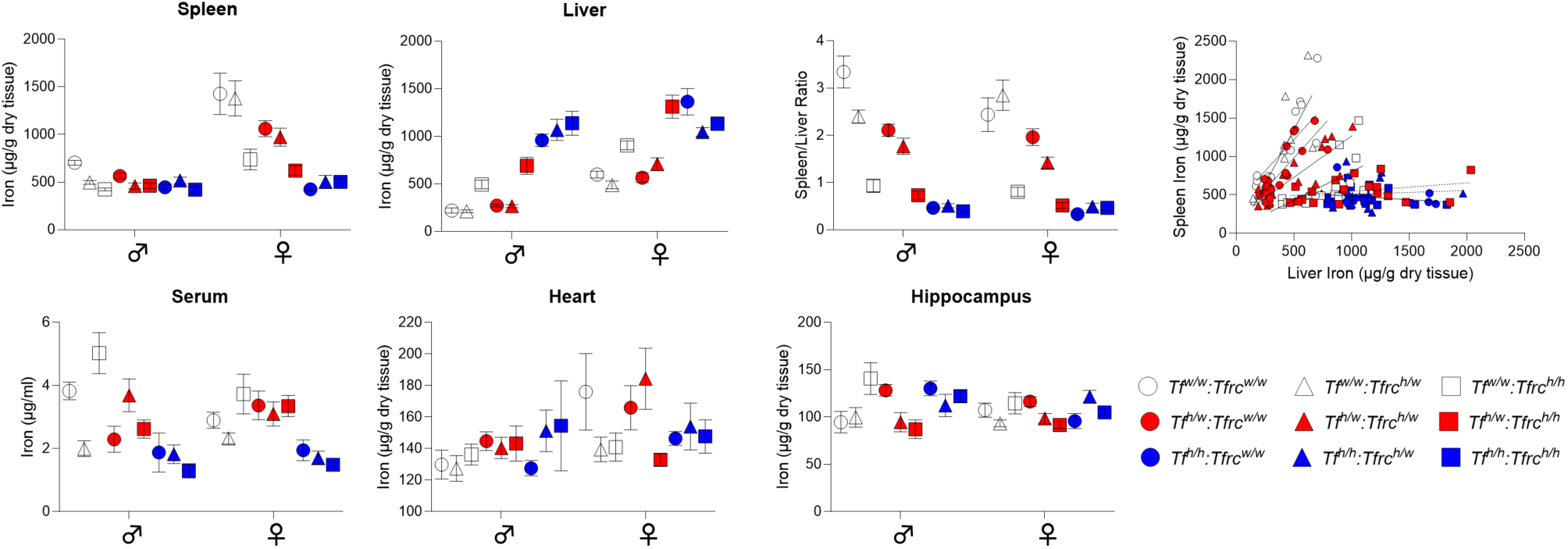
Altered systemic iron distribution in humanized *Tf* and *Tfrc* knock-in rats. Non-heme iron levels in spleen, liver, serum, heart, and hippocampus are shown across genotypes and sexes, along with spleen-to-liver iron ratios and liver–spleen iron regression plots. Two-way ANOVA revealed significant genotype effects in liver and spleen, with altered iron distribution patterns in humanized KI models. Data are represented as mean ± SEM; multiple comparisons are detailed in Table 5, and regression analysis results are summarized in Table 6.

**Table 5.** Statistical analysis of tissue iron levels shown in Figure 17.

| Two-ANOVA summary | Spleen |  | Liver |  | Spleen/Liver |  |
| --- | --- | --- | --- | --- | --- | --- |
| Interaction | F (8, 132) = 7.750<br>p<0.0001 |  | F (8, 132) = 4.195<br>p=0.0002 |  | F (8, 132) = 2.396<br>p=0.0191 |  |
| Sex factor | F (1, 132) = 81.52<br>p<0.0001 |  | F (1, 132) = 75.37<br>p<0.0001 |  | F (1, 132) = 4.353<br>p=0.0389 |  |
| Genotype factor | F (8, 132) = 13.50<br>p<0.0001 |  | F (8, 132) = 40.09<br>p<0.0001 |  | F (8, 132) = 76.27<br>p<0.0001 |  |
| **Tukey's multiple comparisons** | ♂ | ♀ | ♂ | ♀ | ♂ | ♀ |
| $Tf^{w/w}:Tfrc^{w/w}$ vs. $Tf^{w/w}:Tfrc^{h/w}$ | 0.7737 | >0.9999 | >0.9999 | 0.9886 | **0.0037** | 0.7007 |
| $Tf^{w/w}:Tfrc^{w/w}$ vs. $Tf^{w/w}:Tfrc^{h/h}$ | 0.4419 | **<0.0001** | 0.3793 | 0.0888 | **<0.0001** | **<0.0001** |
| $Tf^{w/w}:Tfrc^{w/w}$ vs. $Tf^{h/w}:Tfrc^{w/w}$ | 0.9676 | **0.0365** | >0.9999 | >0.9999 | **<0.0001** | 0.3935 |
| $Tf^{w/w}:Tfrc^{w/w}$ vs. $Tf^{h/w}:Tfrc^{h/w}$ | 0.6028 | **0.0039** | >0.9999 | 0.9823 | **<0.0001** | **0.0003** |
| $Tf^{w/w}:Tfrc^{w/w}$ vs. $Tf^{h/w}:Tfrc^{h/h}$ | 0.6456 | **<0.0001** | **0.0051** | **<0.0001** | **<0.0001** | **<0.0001** |
| $Tf^{w/w}:Tfrc^{w/w}$ vs. $Tf^{h/h}:Tfrc^{w/w}$ | 0.7581 | **<0.0001** | **<0.0001** | **<0.0001** | **<0.0001** | **<0.0001** |
| $Tf^{w/w}:Tfrc^{w/w}$ vs. $Tf^{h/h}:Tfrc^{h/w}$ | 0.8554 | **<0.0001** | **<0.0001** | **0.0017** | **<0.0001** | **<0.0001** |
| $Tf^{w/w}:Tfrc^{w/w}$ vs. $Tf^{h/h}:Tfrc^{h/h}$ | 0.4647 | **<0.0001** | **<0.0001** | **<0.0001** | **<0.0001** | **<0.0001** |
| Two-ANOVA summary | Serum |  | Heart |  | Hippocampus |  |
| Interaction | F (8, 131) = 1.937<br>p=0.0596 |  | F (8, 132) = 1.169<br>p=0.3225 |  | F (8, 130) = 1.487<br>p=0.1681 |  |
| Sex factor | F (1, 131) = 0.098<br>p=0.7544 |  | F (1, 132) = 5.602<br>p=0.0194 |  | F (1, 130) = 2.924<br>p=0.0867 |  |
| Genotype factor | F (8, 131) = 11.49<br>p<0.0001 |  | F (8, 132) = 1.258<br>P=0.2710 |  | F (8, 130) = 4.832<br>p<0.0001 |  |
| **Tukey's multiple comparisons** | ♂ | ♀ | ♂ | ♀ | ♂ | ♀ |
| $Tf^{w/w}:Tfrc^{w/w}$ vs. $Tf^{w/w}:Tfrc^{h/w}$ | 0.1224 | >0.9999 | >0.9999 | 0.6468 | >0.9999 | 0.9799 |
| $Tf^{w/w}:Tfrc^{w/w}$ vs. $Tf^{w/w}:Tfrc^{h/h}$ | 0.9325 | 0.9952 | >0.9999 | 0.5945 | **0.0408** | 0.9997 |
| $Tf^{w/w}:Tfrc^{w/w}$ vs. $Tf^{h/w}:Tfrc^{w/w}$ | 0.3738 | >0.9999 | 0.9980 | 0.9998 | 0.2474 | 0.9981 |
| $Tf^{w/w}:Tfrc^{w/w}$ vs. $Tf^{h/w}:Tfrc^{h/w}$ | >0.9999 | >0.9999 | 0.9999 | >0.9999 | >0.9999 | 0.9987 |
| $Tf^{w/w}:Tfrc^{w/w}$ vs. $Tf^{h/w}:Tfrc^{h/h}$ | 0.9017 | >0.9999 | 0.9994 | 0.3047 | 0.9998 | 0.9370 |
| $Tf^{w/w}:Tfrc^{w/w}$ vs. $Tf^{h/h}:Tfrc^{w/w}$ | 0.3281 | 0.9820 | >0.9999 | 0.8529 | 0.4634 | 0.9943 |
| $Tf^{w/w}:Tfrc^{w/w}$ vs. $Tf^{h/h}:Tfrc^{h/w}$ | 0.0519 | 0.7480 | 0.9789 | 0.9581 | 0.9407 | 0.9687 |
| $Tf^{w/w}:Tfrc^{w/w}$ vs. $Tf^{h/h}:Tfrc^{h/h}$ | **0.0070** | 0.4068 | 0.9674 | 0.8428 | 0.6341 | >0.9999 |

**Table 6.** Regression analysis of spleen-liver tissue iron levels shown in Figure 17.

|  | Slope | 95% CI | R2 | p value | n |
| --- | --- | --- | --- | --- | --- |
| $Tf^{w/w}:Tfrc^{w/w}$ vs. $Tf^{w/w}:Tfrc^{h/w}$ | 1.641 | 0.14 to 3.14 | 0.3719 | 0.0352 | 12 |
| $Tf^{w/w}:Tfrc^{w/w}$ vs. $Tf^{w/w}:Tfrc^{h/h}$ | 2.846 | 1.80 to 3.89 | 0.692 | <0.0001 | 17 |
| $Tf^{w/w}:Tfrc^{w/w}$ vs. $Tf^{h/w}:Tfrc^{w/w}$ | 0.7435 | 0.25 to 1.24 | 0.3873 | 0.0058 | 18 |
| $Tf^{w/w}:Tfrc^{w/w}$ vs. $Tf^{h/w}:Tfrc^{h/w}$ | 1.665 | 1.09 to 2.24 | 0.6755 | <0.0001 | 20 |
| $Tf^{w/w}:Tfrc^{w/w}$ vs. $Tf^{h/w}:Tfrc^{h/h}$ | 1.075 | 0.73 to 1.42 | 0.7321 | <0.0001 | 18 |
| $Tf^{w/w}:Tfrc^{w/w}$ vs. $Tf^{h/h}:Tfrc^{w/w}$ | 0.109 | -0.06 to 0.28 | 0.1038 | 0.1922 | 18 |
| $Tf^{w/w}:Tfrc^{w/w}$ vs. $Tf^{h/h}:Tfrc^{h/w}$ | -0.08958 | -0.35 to 0.17 | 0.05745 | 0.453 | 12 |
| $Tf^{w/w}:Tfrc^{w/w}$ vs. $Tf^{h/h}:Tfrc^{h/h}$ | 0.05192 | -0.26 to 0.36 | 0.007213 | 0.7296 | 19 |
| $Tf^{w/w}:Tfrc^{w/w}$ vs. $Tf^{w/w}:Tfrc^{h/w}$ | -0.08594 | -0.32 to 0.15 | 0.04109 | 0.4515 | 16 |

#### Spleen iron

compared to wild-type controls (*Tf^w/w^:Tfrc^w/w^*), rats carrying humanized *Tf* or *Tfrc* alleles exhibited genotype- and sex-dependent reductions in splenic iron content. These effects were particularly prominent in females. Significant reductions in spleen iron were observed in female *Tf^w/w^:Tfrc^h/h^*, *Tf^h/w^:Tfrc^w/w^*, *Tf^h/w^:Tfrc^h/w^* rats, and in all combinations involving *Tf^h/h^* and/or *Tfrc^h/h^*. In contrast, no significant differences in spleen iron were detected in males, and *Tf^w/w^:Tfrc^h/w^*rats showed no reduction in spleen iron in either sex— making this the only genotype that fully preserved splenic iron homeostasis. These findings indicate that Tf and Tfrc humanization, even in heterozygosity, impairs iron retention in the spleen in a female-specific manner, likely reflecting altered reticuloendothelial iron recycling.

#### Liver iron

Several genotypes exhibited significantly elevated hepatic iron levels in both sexes, including *Tf^w/w^*:*Tfrc^h/h^*, *Tf^h/w^*:*Tfrc^h/h^*, *Tf^h/h^*: *Tfrc^w/w^*, *Tf^h/h^*:*Tfrc^h/w^*, and *Tf^h/h^*:*Tfrc^h/h^*^. No significant changes were observed in simple heterozygotes (*Tf^w/w^*:*Tfrc^h/w^*or *Tf^h/w^*:*Tfrc^w/w^*) or in double heterozygotes (*Tf^h/w^*:*Tfrc^h/w^*). These results indicate that hepatic iron accumulation is a robust phenotype associated primarily with homozygous humanization.

#### Spleen-to-liver iron ratio

Consistent with the combined reduction in spleen iron and increase in liver iron, the spleen/liver iron ratio was significantly reduced in multiple genotypes. In males, a mild but statistically significant reduction in the spleen-to-liver iron ratio was observed in *Tf^w/w^:Tfrc^h/w^*rats (p = 0.0365). More pronounced and highly significant reductions (p < 0.001) were seen in all genotypes carrying either homozygous *Tf^h/h^* or *Tfrc^h/h^* alleles, consistent with a genotype-dependent shift in systemic iron distribution. In females, the spleen-to-liver iron ratio was significantly reduced in all genotypes involving homozygous *Tf^h/h^*or *Tfrc^h/h^* alleles, as well as in double heterozygous *Tf^h/w^:Tfrc^h/w^* rats (all p < 0.001 vs. wild-type). No significant changes were observed in simple heterozygotes (*Tf^w/w^:Tfrc^h/w^* or *Tf^h/w^:Tfrc^w/w^*), indicating that compound heterozygosity or full homozygosity is required to shift iron distribution in females. These data suggest a consistent shift in iron distribution from the spleen toward the liver in humanized animals, particularly in the homozygous state. This altered distribution likely reflects changes in iron recycling dynamics due to partial or complete replacement of the endogenous Tf–TfR1 interaction.

To explore whether iron redistribution disrupted systemic coordination of iron handling, we performed linear regression between spleen and liver iron levels across genotypes. In wild-type and heterozygous humanized animals (*Tf^w/w^:Tfrc^h/w^*, *Tf^h/w^:Tfrc^w/w^*, *Tf^w/w^:Tfrc^h/h^*, *Tf^h/w^*:*Tfrc^h/w^*), spleen and liver iron levels were positively correlated, indicating coordinated regulation. In contrast, in animals with homozygous *Tf* and/or *Tfrc* alleles, this correlation was lost, indicating disruption of systemic iron balance.

#### Serum iron

Serum iron levels remained largely unchanged across genotypes, except for a significant reduction in double homozygous males (*Tf^h/h^*:*Tfrc^h/h^*). This suggests that systemic iron availability is well maintained, even under conditions of tissue redistribution, except in the most severely altered genotype where buffering capacity may be exceeded.

#### Hippocampal iron

CNS iron levels were broadly preserved, except for a modest but significant increase in hippocampal iron in male rats homozygous for *Tfrc* (*Tf*^w/w^:*Tfrc*^h/h^). No changes were detected in females. These findings suggest that while *Tfrc* humanization can affect brain iron content in a sex-specific manner, effects on the CNS are limited.

#### Heart iron

No significant differences were observed in heart iron levels across any genotypes or sexes, indicating that cardiac iron metabolism remains stable following *Tf* or *Tfrc* humanization.

To validate the biochemical quantification of hepatic iron, we performed Prussian blue staining on liver sections from wild type (*Tf^w/w^:Tfrc^w/w^*), double heterozygous (*Tf^h/w^:Tfrc^h/w^*), and *Tf^h/w^:Tfrc^h/h^* rats of both sexes (Figure 18). Wild-type livers showed minimal staining, confirming low iron levels. Double heterozygotes exhibited mild periportal iron deposition, particularly in females, indicating subtle, sex-dependent changes. Strikingly, *Tf^h/w^:Tfrc^h/h^*animals displayed widespread lobular and periportal iron accumulation, with more severe staining in females. This spatial progression parallels phenotypes in models of disrupted iron recycling, such as TfR2 knockouts models (20). Overall, the staining confirms genotype- and sex-dependent hepatic iron accumulation observed in biochemical assays.

**Figure 18.**
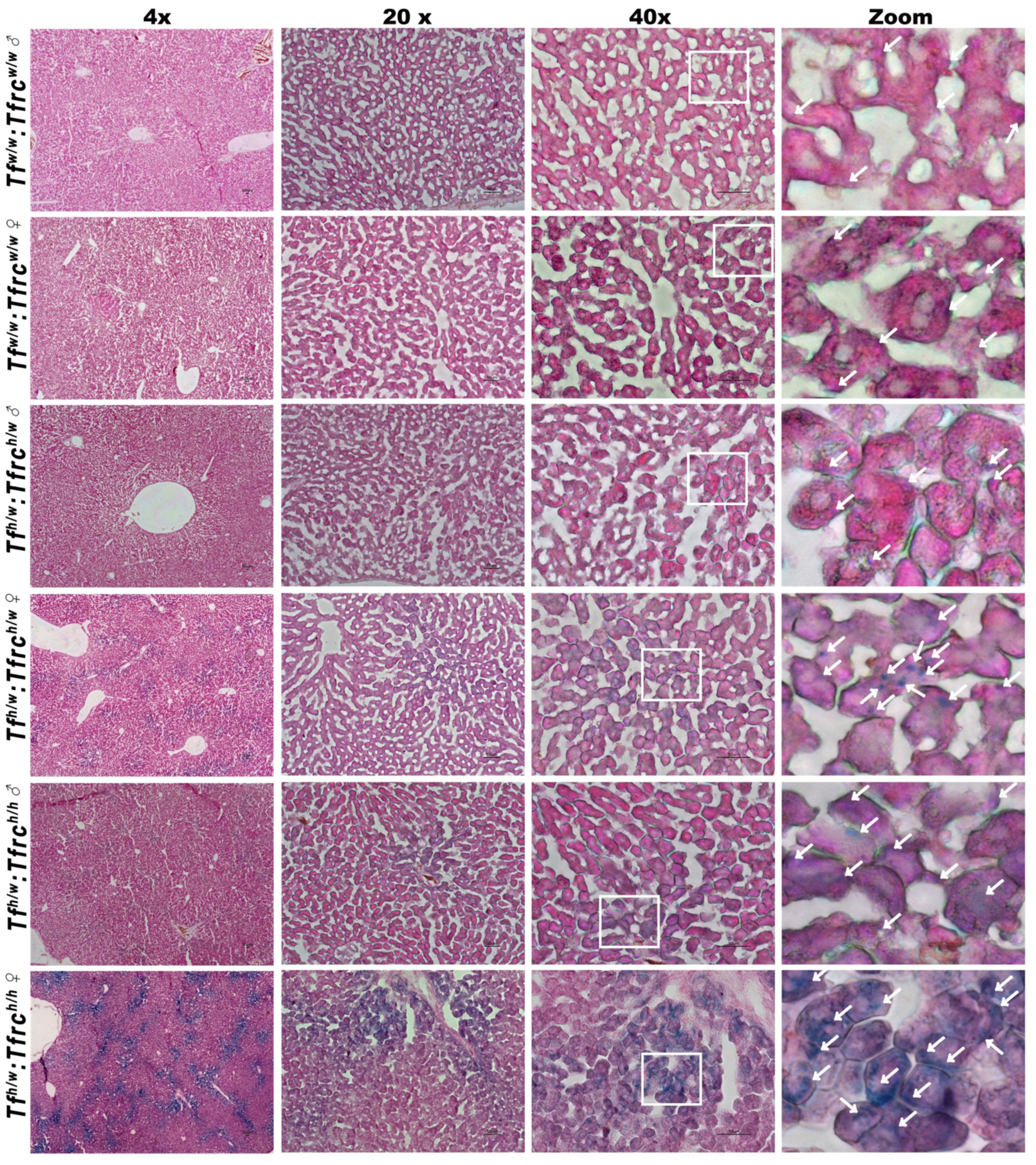
Prussian blue staining of liver sections reveals altered hepatic iron accumulation in *Tf^h/w^:Tfrc^h/h^*rats. Representative images of liver sections stained with Prussian blue to visualize non-heme iron deposition in male and female rats of the following genotypes: *Tf^w/w^:Tfrc^w/w^*, *Tf^h/w^:Tfrc^h/w^*, *Tf^h/w^:Tfrc^h/h^*. Increased staining intensity was observed in *Tf^h/w^:Tfrc^h/h^* animals of both sexes, indicating elevated hepatic iron content.

#### Conclusions

We developed humanized *Tfrc* and *Tf* knock-in rats to enable preclinical evaluation of NewroBus-based therapeutics, which selectively target the human TfR1 axis to cross the BBB. Physiological expression patterns of the human alleles were confirmed via transcript and protein analyses. However, our integrated phenotyping uncovered genotype- and sex-dependent alterations in systemic iron homeostasis, most pronounced in homozygous animals. Despite these abnormalities, even double-homozygous rats were viable, fertile, and able to breed, indicating that the human alleles are sufficiently expressed and functional to compensate for the absence of endogenous rat TfR1 and Tf proteins.

Rats homozygous for humanized *Tf* or *Tfrc* alleles exhibited microcytic, hypochromic anemia with elevated RDW, along with significantly elevated hepatic iron and reduced splenic iron, indicating iron-restricted erythropoiesis and redistribution of iron storage. These phenotypes mirror aspects of human conditions such as hereditary hypotransferrinemia and iron-refractory iron deficiency anemia. Notably, these impairments were more severe in females, aligning with the greater prevalence of iron deficiency in women, potentially due to sex hormones or reproductive iron demands. Given the physiological pattern of expression of the human alleles, this partial functional activity may stem from reduced steady-state protein levels—particularly for human transferrin—and possibly from suboptimal integration into endogenous signaling or trafficking pathways due to species-specific sequence divergence.

In contrast, heterozygous rats maintained largely normal CBC and iron profiles. These animals showed only mild or subclinical changes, such as small reductions in MCV and MCH, and preserved spleen-to-liver iron ratios, suggesting that partial humanization is well tolerated and sufficient for testing human-specific, NewroBus-based therapeutics without overt physiological disruption.

Although homozygous animals present clear impairments in iron handling, they remain useful for in vivo evaluation of NewroBus safety and pharmacodynamics. These genotypes allow for detection of potential interference with the human TF–TfR1 axis—such as impaired iron uptake—without compensation from endogenous rat proteins. Importantly, the viability and fertility of these animals indicate that the human alleles are functionally sufficient to support life, making them a viable, though not ideal, platform to assess possible effects of TfR1-targeting therapeutics on iron homeostasis. Moreover, SDS-PAGE analysis of cerebrospinal fluid confirmed that BBB integrity is preserved across all genotypes, ensuring that these models remain suitable for assessing CNS delivery and safety of BBB-penetrant biologics.

## Experimental Procedures

### Animals

All animal procedures were conducted in accordance with the NIH *Ethical Guidelines for the Treatment of Laboratory Animals*. All protocols were approved by the Rutgers Institutional Animal Care and Use Committee (IACUC Protocol #201702513). Efforts were made to minimize animal suffering and reduce the number of animals used.

### Generation of humanized *Tfrc* and *Tf* Long Evans Rats

The generation of humanized rats was outsourced to Cyagen. Complete details of the targeting strategy, construct design, and validation data are provided in the *Supporting Information*. A summary is included here.

#### Humanized *Tfrc* rats

The *Tfrc* gene (NCBI RefSeq: NM_022712.1) is located on rat chromosome 11 and contains 19 exons, with the ATG start codon in exon 2 and the TAA stop codon in exon 19. The corresponding human gene, *TFRC* (NCBI RefSeq: NM_001128148.3), also contains 19 exons (Transcript: ENST00000360110), with the same exon organization.

For the knock-in model, a portion of rat exon 2 (32 bp) was replaced with a cassette containing the human *TFRC* coding sequence, followed by the rat *Tfrc* 3′ UTR and an SV40 polyadenylation signal (*“Human TFRC CDS–3′UTR of rat Tfrc–3×SV40 pA”*). The CRISPR-Cas9 system was used for genome editing, with co-injection of Cas9 mRNA, guide RNAs (gRNAs), and the targeting vector into fertilized rat embryos.

#### Humanized *Tf* rats

The rat *Tf* gene (NCBI RefSeq: NM_001013110.1) is located on chromosome 8 and comprises 17 exons, with the ATG start codon in exon 1 and the TAA stop codon in exon 17. The human *TF* gene (NCBI RefSeq: NM_001063.4), located on human chromosome 3, also contains 17 exons with the same exon-intron structure (Transcript: ENST00000402696).

For the knock-in model, a portion of exon 2 of the rat *Tf* gene (140 bp) was replaced with a cassette containing the human *TF* coding sequence (amino acids 20–698), followed by the rat *Tf* 3′ UTR and an SV40 polyadenylation signal (*“Human TF CDS(aa 20–698)–3′UTR of rat Tf–3×SV40 pA”*). The endogenous rat signal peptide (amino acids 1–19) was retained to ensure proper protein processing and secretion in the rat background.

To generate the targeting vector, 5′ and 3′ homology arms were amplified from rat genomic DNA using PCR. Genome editing was performed by co-injecting Cas9 mRNA, guide RNAs (gRNAs), and the targeting vector into fertilized rat embryos.

### Protein preparation

These procedures were performed as previously described (21). Briefly, the rats were first put under anesthesia using isoflurane, followed by perfusion through intracardiac catheterization using ice-cold PBS. Brains were extracted and homogenized with a glass-Teflon homogenizer in 250 mM Sucrose, 20 mM Tris-base pH 7.4, 1 mM EDTA, 1 mM EGTA plus protease and phosphatase inhibitors (Thermo Scientific). All steps were carried out on the ice. Homogenates were sonicated at 20s at 50% amplitude with 30s interval resting every 10s sonication. The protein content of sonicated homogenates was quantified using the Bradford method.

### Tissue and CSF collection

These procedures were performed as previously described (12). Briefly, the rats were first put under anesthesia using isoflurane, followed by perfusion through intracardiac catheterization using ice-cold PBS. Blood was collected via cardiac puncture before perfusion, drawn into serum separator tubes (BD Becton Dickinson vacutainers, SSTTM) and left to incubate at room temperature for 30 minutes. The tubes were then centrifuged at 2000×g for 10 minutes to separate the serum, which was subsequently stored at -80°C. CSF was rapidly collected after perfusion and saved in liquid nitrogen. Spleen, Liver, Lung, Heart and Brain were dissected post PBS perfusion. The brain was further separate into hippocampus and cortex. Each tissue was collected as two similar size pieces and saved into two independent tubes. One piece of tissue was weighed for ion measurement and the other piece was immediately frozen in liquid nitrogen.

### RNA isolation, RT and qRT-pcr analysis

Rat tissues were extracted from 3-month-old rats after rapidly intracardiac ice-cold PBS perfusion. Spleen, Liver, Heart, Kidney and Brain were rapidly dissected post PBS perfusion and homogenized with a glass-teflon homogenizer in 250 mM Sucrose, 20 mM Tris-base pH 7.4, 1 mM EDTA, 1 mM EGTA plus protease and phosphatase inhibitors. 50µl of homogenate was added with 300ul RLT buffer (Qiagen RNeasy kit 74106) w/w 1% β-mercaptoethanol and rapidly frozen in liquid nitrogen and later used for RNA extraction. Total RNA was extracted with RNeasy RNA Isolation kit (Qiagen 74106) and used to generate cDNA with a High-Capacity cDNA Reverse Transcription Kit (Thermo 4368814) with random hexamer priming. Real time polymerase chain reaction was carried out with TaqMan Fast Advanced Master Mix (Thermo 4444556), and the appropriate TaqMan (Thermo) probes. Reference gene *Gapdh* was detected with probe Rn01775763_g1 and target genes were detected with Hs00169070_m1 (*TF*, human Transferrin), Rn01445482_m1 (*Tf,* rat Transferrin), Hs00951083_m1 (*TFRC*, human Transferrin Receptor1) and Rn01474701_m1 (*Tfrc*, rat Transferrin receptor 1). Samples were analyzed on a QuantStudio 6 Flex Real-Time PCR System (Thermo 4485697), and relative RNA amounts were quantified using LinRegPCR software (hartfaalcentrum.nl).

### Immunoprecipitation

Sonicated homogenate was mixed with 10% protein A/G beads (Thermo, 20421) and rotated for 2 hours at 4°C to avoid non-specific beads related binding. Homogenate after pre-cleaning was used as input for immunoprecipitation. Rat specific Tfr1 antibody (Thermo, MA170033) was used to immunoprecipitate at 1:100 dilution. Immunocomplexes were washed 5 times with homogenate buffer and proteins bound were eluted with 2× LDS sample buffer with (reduced) or without (non-reduced) 20% β-mercaptoethanol (Invitrogen; NP0007) at 95 °C. Input (total lysates, T.L.) and immunoprecipitation (I.P.) eluates were analyzed by Western blot.

### Western blot (WB)

WB were performed as follows: proteins were diluted with Homogenate buffer and LDS Sample Buffer (Invitrogen NP0007) containing 20% β-mercaptoethanol to a final concentration of 1 μg/μl. For non-reducing conditions, samples were prepared without β-mercaptoethanol. Samples were loaded onto a 4–12% Bis-Tris polyacrylamide gel (Biorad 3450125), running at constant voltage of 130V for 45min and transferred onto nitrocellulose membranes at 25 V for 7 minutes using the Trans-Blot Turbo system (Biorad). Blotting efficiency was confirmed by red Ponceau staining of the membranes.

Membranes were blocked for 45 minutes in 5% nonfat-milk (Biorad 1706404), followed by extensive washing in PBS/0.05% Tween-20. Primary antibodies (anti-TFRC Rabbit mAb, Cell Signaling Technology, 13113; anti-Tfrc Rabbit mAb, Cell Signaling Technology, 55487; anti-TF Rabbit mAb, Cell Signaling Technology, 35293; anti-Tf Rabbit mAb, Invitrogen, MAB529871; anti-GAPDH Rabbit mAb, Sigma, g9545) were used at 1:1000 dilution overnight at 4°C. After three 10 minutes washes with PBS/0.05% Tween-20, membranes were incubated with a mixture of HRP-conjugated anti-rabbit secondary antibodies (Southern Biotech, OB405005 and Cell Signaling Technology, 7074) diluted 1:1,000 in 5% nonfat-milk for 45 minutes at room temperature with shaking. Blots were developed using Clarity Western ECL reagent (Bio-rad 1705061) and visualized on a ChemiDoc MP Imaging System (Bio-Rad). For WB quantifications, signal intensities were analyzed with Image Lab software (Bio-Rad).

### ELISA

To measure the soluble TFRC and TF concentration, the following commercial ELISA kits were used as it suggested: human Transferrin ELISA (Abcam ab187391) and soluble TFR1 ELISA (Meso Scale Discovery K151P9K). Collected CSF and serum were diluted as recommended in assay-specific sample dilution buffer and performed following instructive protocols.

### Immunofluorescence (IF)

Rat tissues including kidney, lung, duodenum, and liver were isolated after transcardiac perfusion with PBS (without calcium and magnesium) followed by 4% paraformaldehyde (PFA) (Electron Microscopy Sciences, Cat. No. 15714-S) at a rate of 10 ml/min. The tissues were transferred to 4% PFA for 24 hours at 4°C on an orbital shaker. After fixation, the tissues were washed twice with PBS, moved into a 30% sucrose solution at 4°C on an orbital shaker for 48 hours, and then embedded with OCT (Fisher, Cat. No. 23-730-571) and stored at -80°C. Sections were cut using a rotary cryostat (Leica CM1950), mounted on charged glass slides (Fisher, Cat. No. 22-037-246) and stored at -80°C. The duodenum and kidney sections were cut at 15µm and the liver, lung and brain sections were cut at 20µm. Prior to staining, slides were brought to room temeperature for 5 minutes, and the barrier was drawn with Super PAP Pen liquid blocker, further rewet with PBS for 10 minutes and blocked for 1 hour at room temeperature with 10% serum containing 0.3% Triton X-100 (Sigma, Cat. No. T9284). Sections were then incubated with the primary antibody (anti-TFRC Rabbit mAb, Cell Signaling Technology, 13113; anti-Tfrc mouse Ab, Cell Thermo, MA170033; anti-TF Rabbit mAb, Cell Signaling Technology, 35293; anti-Tf Rabbit mAb, Invitrogen, MAB529871 diluted at 1:250 in PBS containing 5% serum with 0.3% Triton X-100 overnight at 4°C. The next day, slides were washed three times for 10 minutes each with PBS containing 0.5% serum and 0.3% Triton X-100, then incubated with the secondary antibody (Goat anti-Rabbit IgG (H+L) Cross-Adsorbed Secondary Antibody with Alexa Fluor™ 594, Invitrogen, A11012 and Goat anti-Mouse IgG2a Cross-Adsorbed Secondary Antibody with Alexa Fluor™ 488, Invitrogen, A21132) at 1:1000 in PBS containing 5% serum with 0.3% Triton X-100 for 2 hours at room temeperature, and washed three times with PBS containing 0.3% Triton X-100 at RT. Finally, all IF tissue sections were washed with PBS 2 times for 10 minutes each and mounted with coverslips (Corning, Cat. No. 2980-225) in an aqueous mounting medium containing DAPI (Southern Biotech, Cat. No. 0100-20). Images were captured using a Nikon A1R inverted confocal microscope.

### Prussian blue staining

Tissue collection and OCT embedding were performed as described in the immunohistochemistry section. For iron staining of OCT-embedded tissue sections (10 µm thick, in vitro), we used a commercially available iron stain kit (StatLab, Cat. No. KTIRO) to detect ferric ion levels in rat tissues. Prior to staining, tissue sections were equilibrated to room room temeperature for 5 minutes. A hydrophobic barrier was drawn around each section using a Super PAP Pen liquid blocker, followed by rehydration in PBS for 10 minutes. Sections were then incubated for 20 minutes in a 1:1 solution of 3% hydrochloric acid and 3% potassium ferrocyanide. After incubation, the slides were rinsed in PBS three times, 1 minute each. Nuclear Fast Red stain was applied for 5 minutes to counterstain, followed by another three 1-minute PBS washes. Finally, slides were mounted using a water-soluble mounting medium, Fluoromount-G (Southern Biotech, Cat. No. 0100-01), and brightfield images were captured using the Keyence BZ-X710 microscope.

### CBC measurement

Blood samples were collected from rats via the tail vein using a 25G VACUETTE Safety Blood Collection Set (Greiner Bio-One). The collected blood was immediately transferred into MiniCollect K2EDTA tubes (Greiner Bio-One) to prevent coagulation. Approximately 200 µL of blood was obtained per animal in the morning to minimize circadian variations. After collection, samples were maintained at 4°C until measurement. CBC parameters were measured using the Heska Element HT5 CBC Analyzer. The following hematological parameters were assessed: white blood cell count (WBC, 10^3^/µL), neutrophil count (NEU, 10^3^/µL), lymphocyte count (LYM, 10^3^/µL), monocyte count (MONO, 10^3^/µL), eosinophil count (EOS, 10^3^/µL), basophil count (BAS, 10^3^/µL), neutrophil percentage (NEU%), lymphocyte percentage (LYM%), monocyte percentage (MONO%), eosinophil percentage (EOS%), basophil percentage (BAS%), red blood cell count (RBC, 10^6^/µL), hemoglobin (HGB, g/dL), hematocrit (HCT, %), mean corpuscular volume (MCV, fL), mean corpuscular hemoglobin (MCH, pg), mean corpuscular hemoglobin concentration (MCHC, g/dL), red cell distribution width percentage (RDW%), platelet count (PLT, 10^3^/µL), and mean platelet volume (MPV, fL).

### Tissue iron measurement

Non-heme iron content in tissue samples was quantified using a bathophenanthroline-based colorimetric assay, following previously published methods (22). Tissue samples were first dried at 65°C for 48 hours, after which 10–100 mg of dried tissue was accurately weighed and transferred into 1.5 mL microcentrifuge tubes. Each sample was then incubated with 1 mL of acid mixture (37% hydrochloric acid and trichloroacetic acid) at 65°C for 20 hours to release non-heme iron. Following digestion, 500 µL of the clear supernatant was carefully transferred into a new tube and frozen for batch analysis. For serum iron measurements, blood was collected using a 25G VACUETTE Safety Blood Collection Set (Greiner Bio-One) and transferred into Microtainer SST tubes (BD). Samples were kept at room temperature for 30 minutes, then centrifuged at 9,000 x g for 90 seconds, and the serum was collected and stored at -80°C. For iron quantification, samples were mixed with a working chromogen reagent containing 4,7-diphenyl-1,10-phenanthroline disulfonic acid disodium salt, thioglycolic acid, and sodium acetate buffer in a 96-well microplate. The reaction was allowed to develop at room temperature for 15 minutes, and absorbance was subsequently measured at 535 nm using a microplate reader. A working iron standard solution (11.169 µg Fe/mL) was prepared from carbonyl iron powder, and absorbance values were used to calculate tissue iron content. Final iron concentrations were expressed as µg of iron per gram of dry tissue, with adjustments made for sample volume and reaction conditions.

## Supporting Information

## 1. Extended Experimental procedures

### 1.1 generation of TFRC knock-in rat

The KI founder F0-*Tfrc^h^* rat was generated by CRISPR/Cas-mediated genome engineering. The rat *Tfrc* gene (GenBank accession number: NM_022712.1) is located on rat chromosome 11. It comprises 19 exons, with ATG start codon in exon 2 and TAA stop codon in exon 19. The human TFRC gene (NCBI Reference Sequence: NM_001128148.3) is located on human chromosome 3. 19 exons have been identified, with the ATG start codon in exon 2 and the TAA stop codon in exon 19(Transcript: 201-ENST00000360110). For the KI model, partial exon 2 (32bp) of rat Tfrc will be replaced with the “Human TFRC CDS-3’UTR of rat Tfrc-3*SV40 pA” cassette. One pair of gRNA targeting vectors were designed as following,

gRNA-A1 (matches forward strand of gene): CAGATCAGCATTCTCTAACT-TGG

gRNA-B1 (matches forward strand of gene): CAGCATTCTCTAACTTGGTA-AGG.

*Off-target analysis of targeting sequence* gRNA: Out of all potential gRNAs, four have been selected for their position and a low number of predicted off-targets. The off-target analysis is based on the Rat (Rattus norvegicus) Ensembl V 103. Predicted off-target sites with up to one mismatch in the seed and three or four total mismatches are listed. Lowercase letters indicate mismatches to the target site. (Legend for off-target site positon: **E** = exonic; **I** = intronic; **-** = intergenic; Legend for the CRISPRater score: LOW efficacy (score<0.56); MEDIUM efficacy (0.56<=score<=0.74); HIGH efficacy (score>0.74)) Off-target analysis for gRNA-A1: Sequence: CAGATCAGCATTCTCTAACTTGG

Efficacy score by CRISPRater: 0.61 MEDIUM

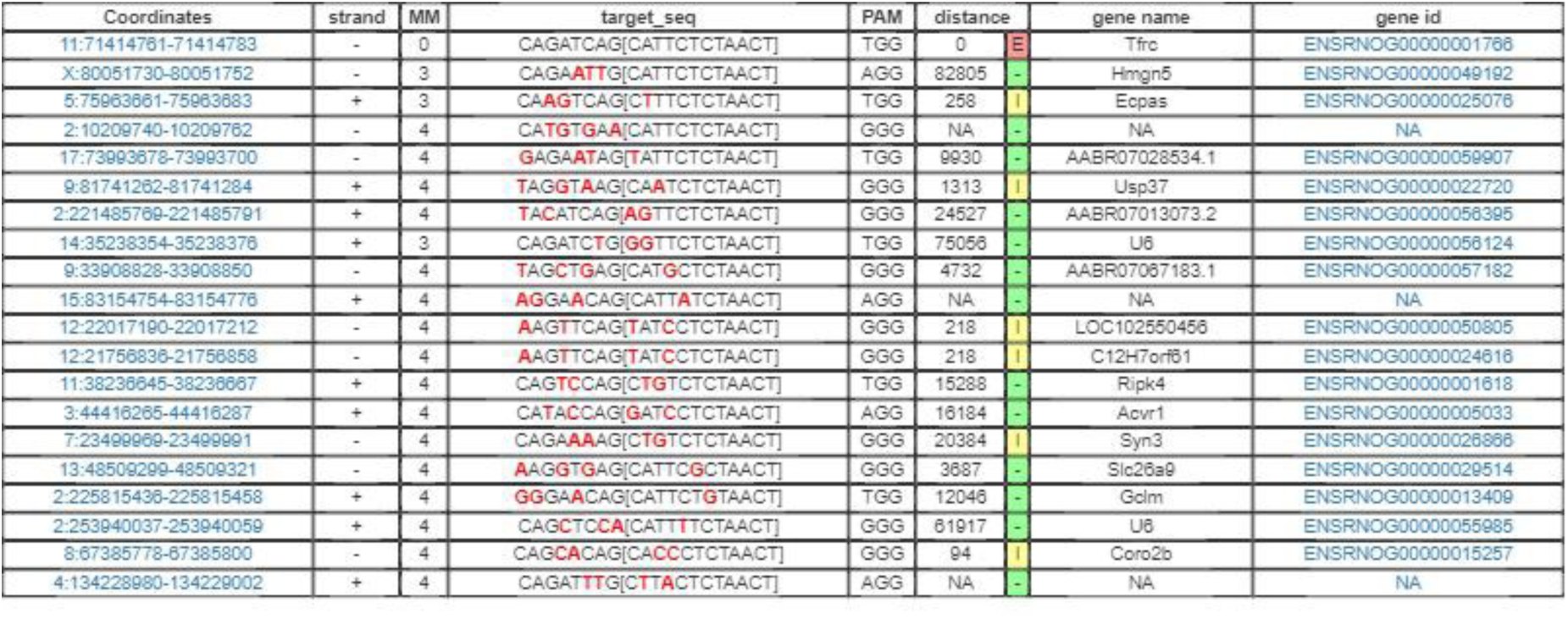

Off-target analysis for gRNA-B1: Sequence: CAGCATTCTCTAACTTGGTAAGG

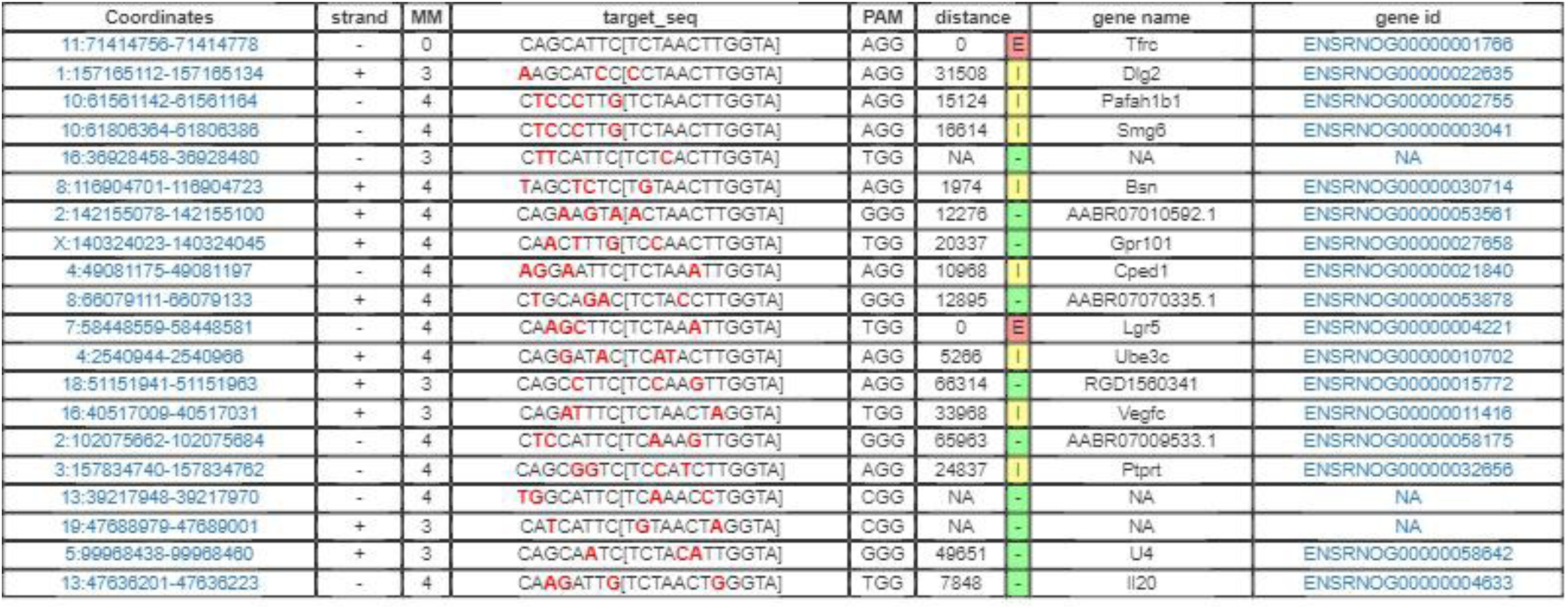

Efficacy score by CRISPRater: 0.61 MEDIUM

Target Vector building: Mouse genomic fragments containing homology arms (HAs) were amplified from BAC clone by using high fidelity Taq DNA polymerase, and were sequentially assembled into a targeting vector together with recombination sites and selection markers shown below.

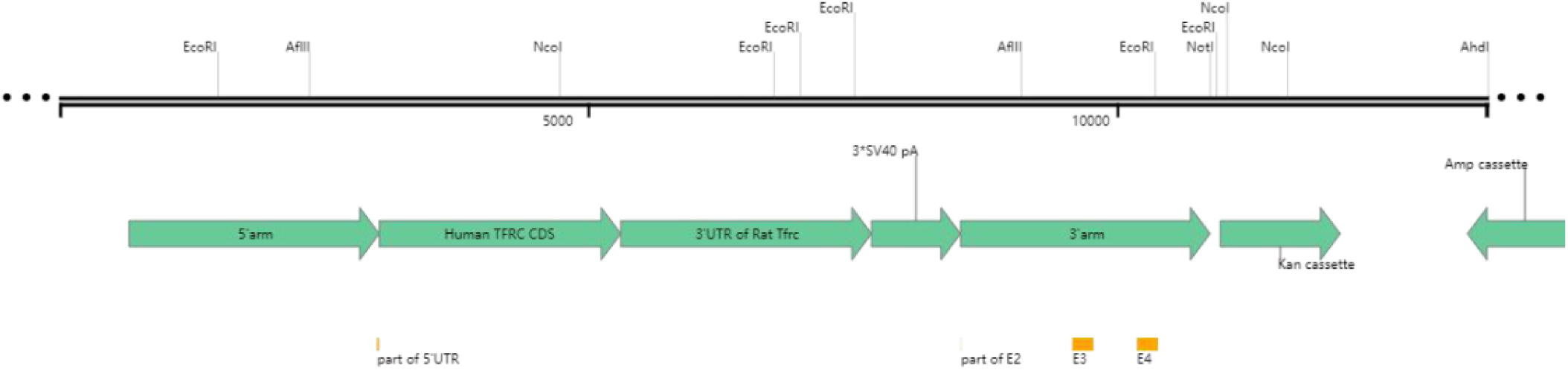

Targeting vector was digested by restriction enzymes for confirmation purposes. Units below left are all in kilo-base pair (kb). (1.AhdI+NcoI: 6.3/5.6/1.9/0.6; 2. AflII+EcoRI: 5.0/4.4/1.6/1.3/0.9/0.6/0.5/0.2; 3.NotI: 14.4). Vectors were further confirmed by sequencing.

The gRNA to rat Tfrc gene, the donor vector containing “Human TFRC CDS-3’UTR of rat Tfrc-3*SV40 pA” cassette, and Cas9 mRNA were co-injected into fertilized rat eggs to generate targeted knockin offspring. F0 founder animals were identified by PCR followed by sequence analysis, which were bred to wildtype rats to test germline transmission and F1 animal generation. The rat *Tfrc* locus was amplified by PCR with the following 2 pairs of specific forward (F) and reverse (R) primers: F1-ATATTGGAACACTTGTGAGGGTG, R1-TCCCATAGCAGATACTTCCACTAC and F2-CCGGATCAGCTTGATGGGGAT, R2-AAAGCCTCATCCTTACATATCTGC.

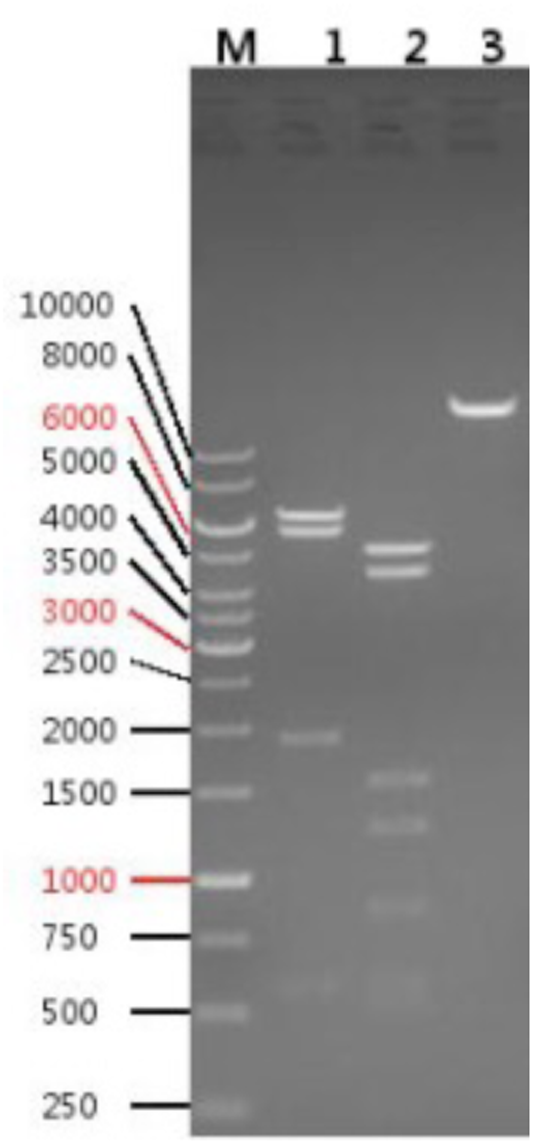

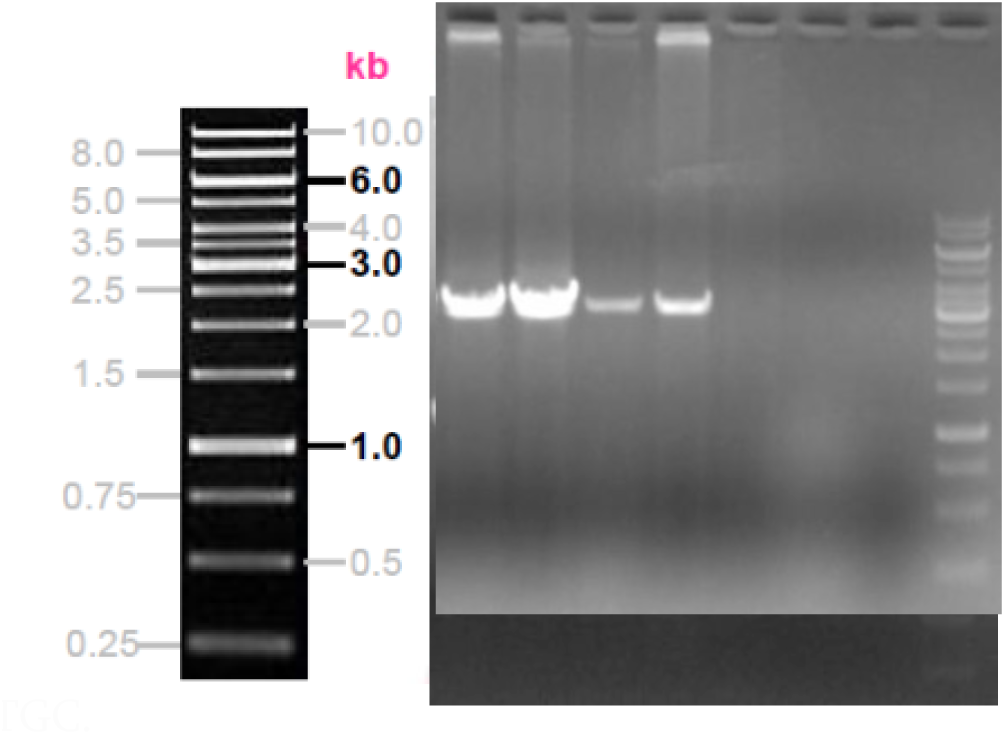

4 pups (2 males and 2 females) with ID 2,9,16 and 26 were selected as positive by PCR screening (see right, top was using F1/R1 pair of primers and bottom was using F2/R2 pair of primers).

Cas9 RNA, sgRNA and oligo donor are co-injected into zygotes, but homology-directed repair can occur even after few cell cycles. Thus, injected rats can have a mixture of correctly targeted alleles and alleles carrying aberrant mutations or no mutations. To identify rats carrying correctly targeted *TFRC* alleles, the PCR products were cloned into TA vectors and were sequenced using forward primer F-GCCAGCCAGAACTTCCTAGTAGAT and F-CCGGATCAGCTTGATGGGGAT.

Southern blot confirmation. The correct gene targeting in 4 F1 animals (2, 9, 16 and 26) were confirmed by Southern blot analysis of the tail DNA samples. The strategy of Southern blot analysis is shown below.

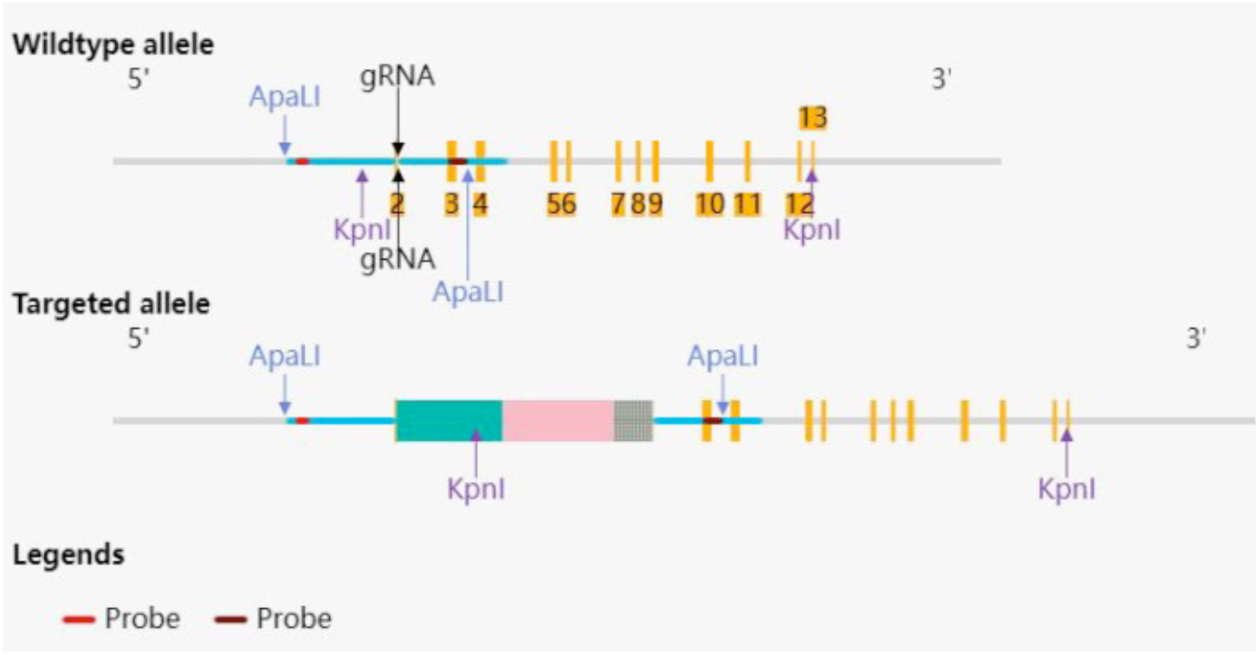

Expected Fragment Sizes for Southern Blot:

5’Probe-ApaLI: 3.92 kb-WT, 9.38 kb-MT

3’Probe-KpnI: 9.63 kb-WT, 12.66 kb-MT

Primers for 5’Probe:

F-CTTCCTTCTGTGTGAACGCAGGAT

R-AGATGTCACCAGTGCCAACGTCTGT

Primers for 3’Probe:

F-GAGCCATTGTCATACACCCGGTTT

R-TGCACACTTTGCCCTAATGGGAG-

All of the four rats (2, 9, 16 and 26) were confirmed correct by Southern blot analysis.

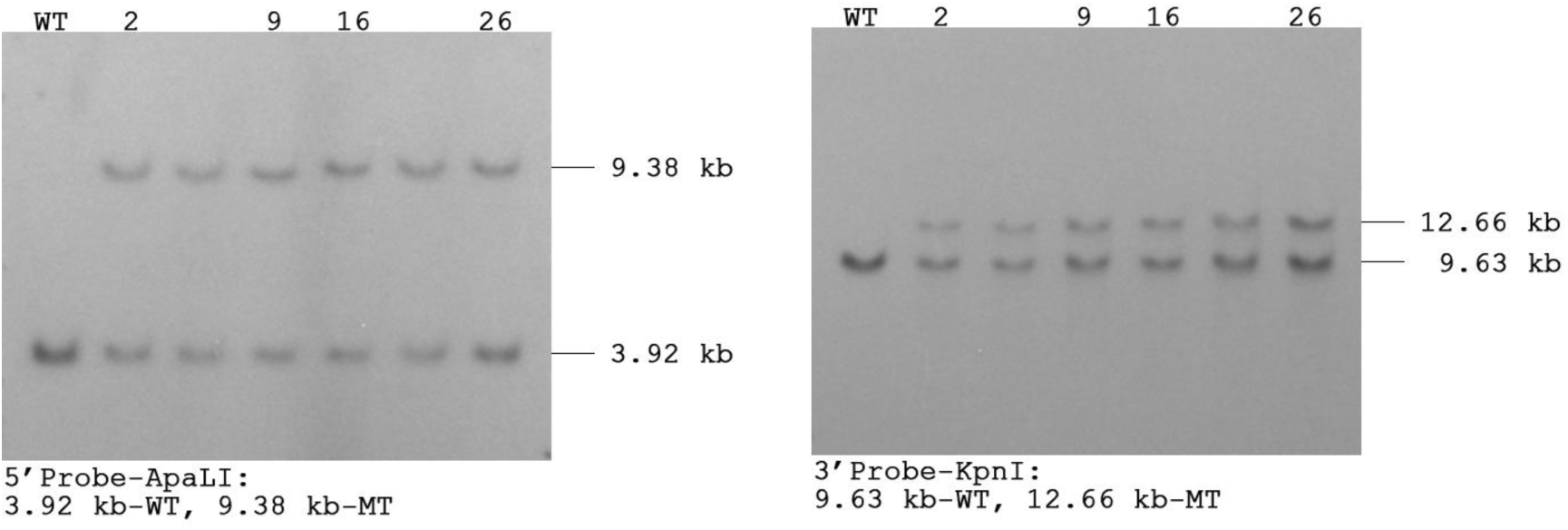

### 1.2 generation of TF knock-in rat

The KI founder F0-*Tf^h^* rat was generated by CRISPR/Cas-mediated genome engineering. The rat *Tf* gene (GenBank accession number: NM_001013110.1) is located on rat chromosome 8. It comprises 17 exons, with ATG start codon in exon 1 and TAA stop codon in exon 17. The human *TF* gene (NCBI Reference Sequence: NM_001063.4) is located on human chromosome 3. 17 exons have been identified, with the ATG start codon in exon 1 and the TAA stop codon in exon 17 (Transcript: 201-ENST00000402696). For the KI model, partial exon 2 (140bp) of rat Tf will be replaced with the “Human TFCDS(aa.20∼698)-3’UTR of rat Tf-3*SV40pA” cassette. Two pairs of gRNA targeting vectors were designed as following,

gRNA-A1 (matches reverse strand of gene): ACCATTTGACCGTTTTGTCA-GGG

gRNA-A2 (matches reverse strand of gene): CTTGATGCAATCTTGATAGG-AGG

gRNA-B1 (matches reverse strand of gene): CACCATTTGACCGTTTTGTC-AGG

gRNA-B2 (matches reverse strand of gene): CTCCTATCAAGATTGCATCA-AGG

*Off-target analysis of targeting sequence* gRNA: Out of all potential gRNAs, four have been selected for their position and a low number of predicted off-targets. The off-target analysis is based on the Rat (Rattus norvegicus) Ensembl V 103. Predicted off-target sites with up to one mismatch in the seed and three or four total mismatches are listed. Lowercase letters indicate mismatches to the target site. (Legend for off-target site positon: **E** = exonic; **I** = intronic; **-** = intergenic; Legend for the CRISPRater score: LOW efficacy (score<0.56); MEDIUM efficacy (0.56<=score<=0.74); HIGH efficacy (score>0.74))

Off-target analysis for gRNA-A1: Sequence: ACCATTTGACCGTTTTGTCA-GGG

Efficacy score by CRISPRater: 0.69 MEDIUM

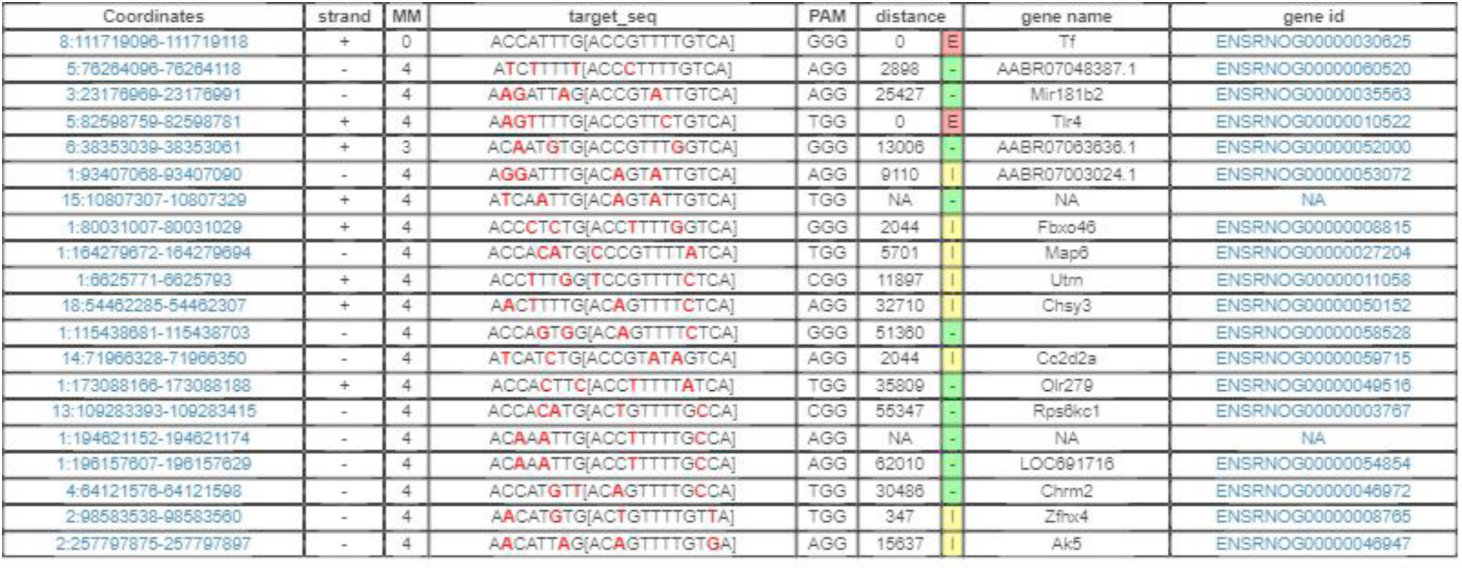

Off-target analysis for gRNA-A2: Sequence: CTTGATGCAATCTTGATAGG-AGG

Efficacy score by CRISPRater: 0.70 MEDIUM

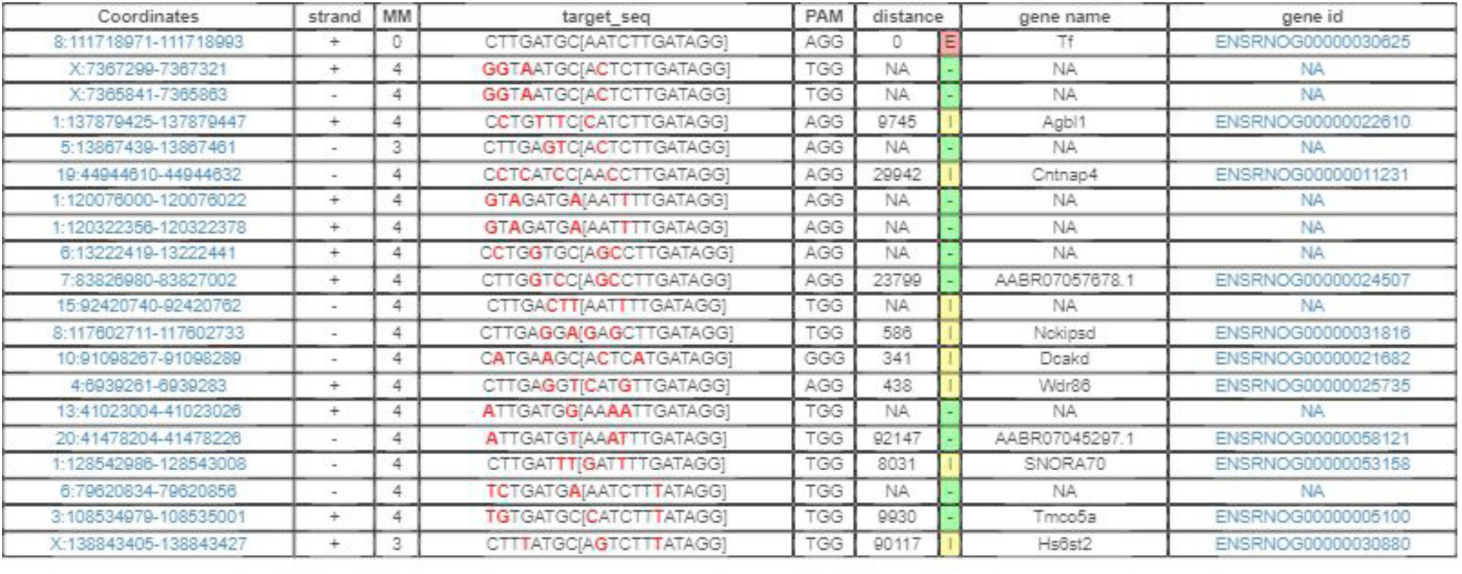

Off-target analysis for gRNA-B1: Sequence: CACCATTTGACCGTTTTGTC-AGG

Efficacy score by CRISPRater: 0.60 MEDIUM

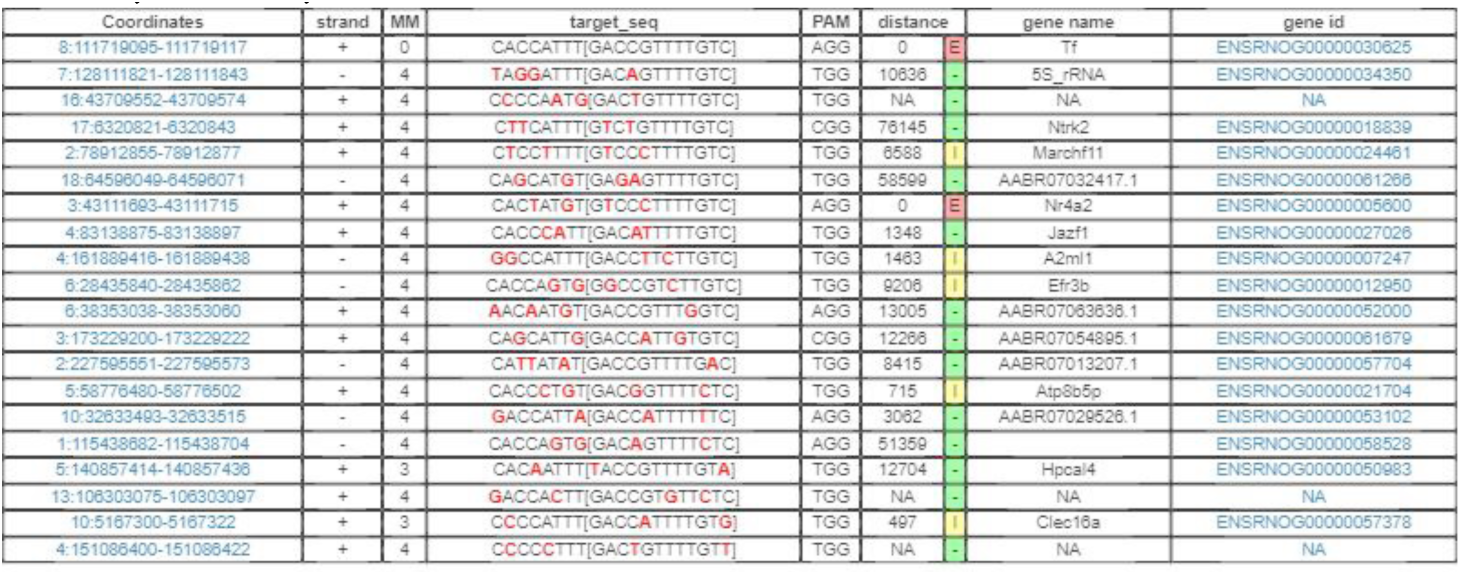

Off-target analysis for gRNA-B2: Sequence: CTCCTATCAAGATTGCATCA-AGG

Efficacy score by CRISPRater: 0.73 MEDIUM

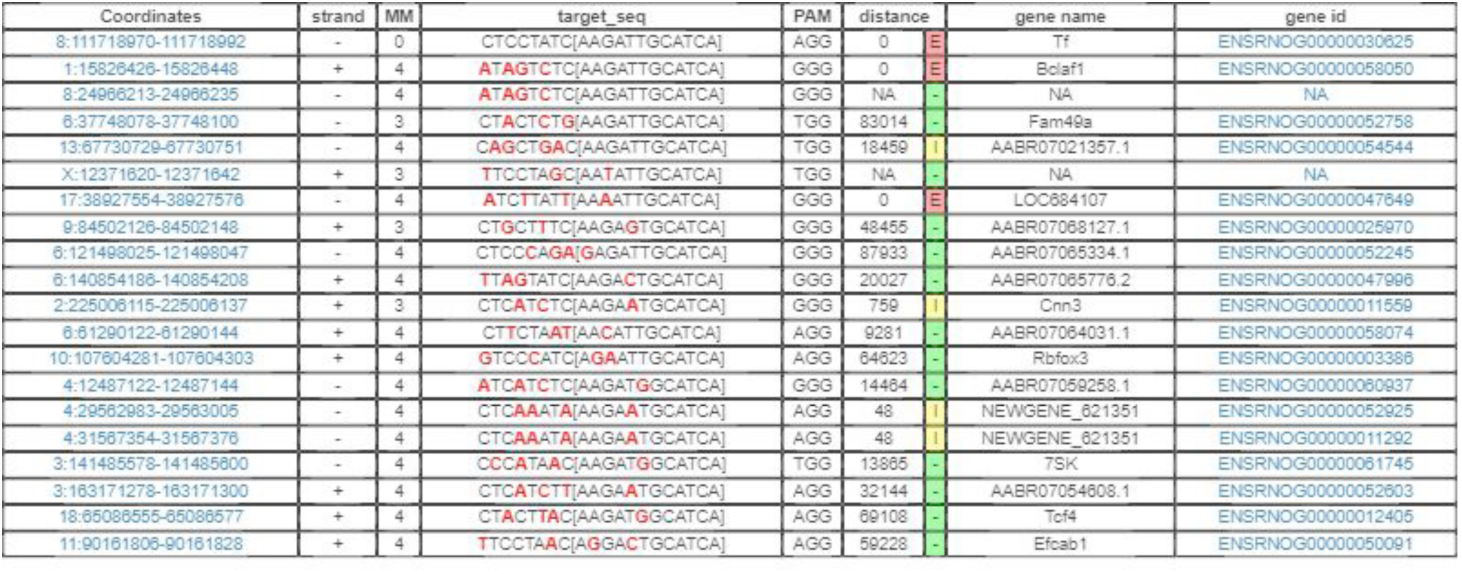

Target Vector building: Mouse genomic fragments containing homology arms (HAs) were amplified from BAC clone by using high fidelity Taq DNA polymerase, and were sequentially assembled into a targeting vector together with recombination sites and selection markers shown below.

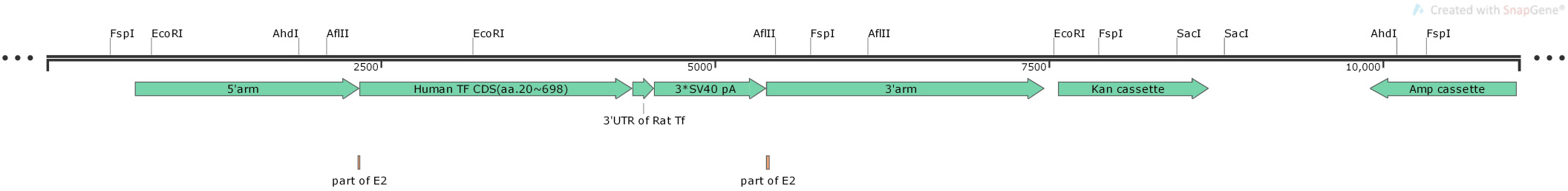

Targeting vector was digested by restriction enzymes for confirmation purposes. Units below left are all in kilo-base pair (kb). (1.FspI: 5.2/2.4/2.2/1.2; 2.AflII+SacI: 4.3/3.4/2.3/0.7/0.4; 3.AhdI+EcoRI: 4.3/2.6/1.7/1.3/1.1). Vectors were further confirmed by sequencing.

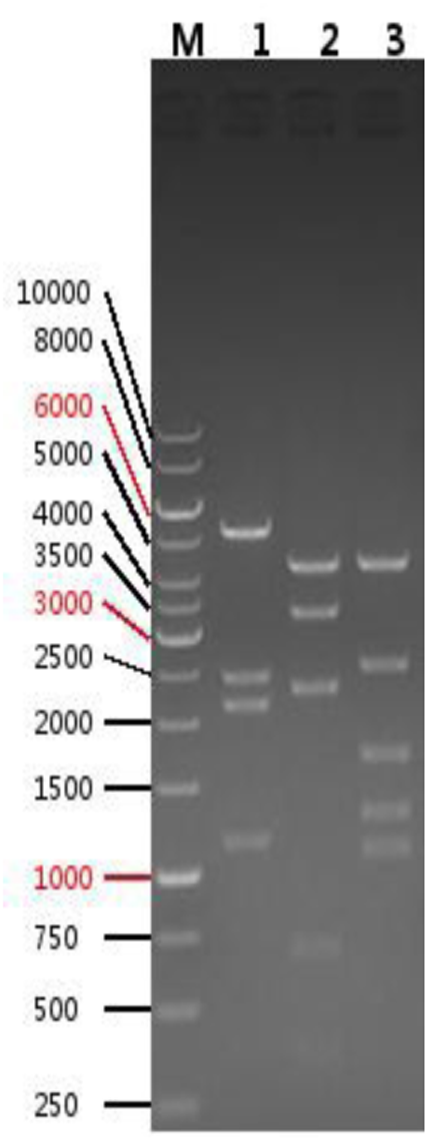

The gRNA to rat *Tf* gene, the donor vector containing “ Human TF CDS-3’UTR of rat Tf-3*SV40pA “ cassette, and Cas9 mRNA were coinjected into fertilized rat eggs to generate targeted knockin offspring. F0 founder animals were identified by PCR followed by sequence analysis, which were bred to wildtype rats to test germline transmission and F1 animal generation. The rat *Tf* locus was amplified by PCR with the following 2 pairs of specific forward (F) and reverse (R) primers:

F1- TTTATTATCCTCCTCACACTATCTC

R1- TGCAGGAATTAGCTTGCAGATC

F2- GAAGGATAGTGGCTTCCAGATGA

R2- GATTTGGTAGACGCAGACAGGAG.

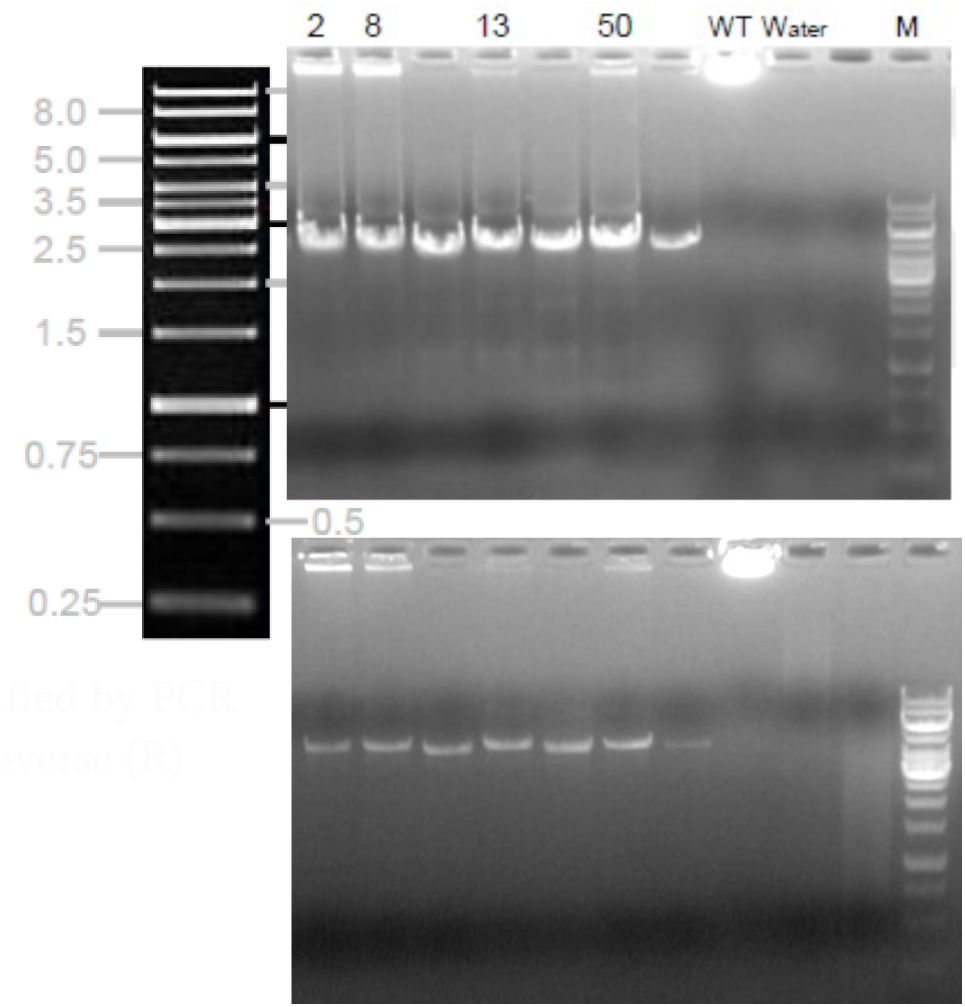

4 pups (2 males and 2 females) with ID 2,8,13 and 50 were selected as positive by PCR screening (see right, top was using F1/R1 pair of primers and bottom was using F2/R2 pair of primers).

Cas9 RNA, sgRNA and oligo donor are co-injected into zygotes, but homology-directed repair can occur even after few cell cycles. Thus, injected rats can have a mixture of correctly targeted alleles and alleles carrying aberrant mutations or no mutations. To identify rats carrying correctly targeted *TFRC* alleles, the PCR products were cloned into TA vectors and were sequenced using forward and reverse primer F-AGTATCAGGGTGCTGGAGAAC and R-ATGCCACAGACATTCCTCTTTCC.

Southern blot confirmation. The correct gene targeting in 4 F1 animals (2, 8, 13 and 50) were confirmed by Southern blot analysis of the tail DNA samples. The strategy of Southern blot analysis is shown below.

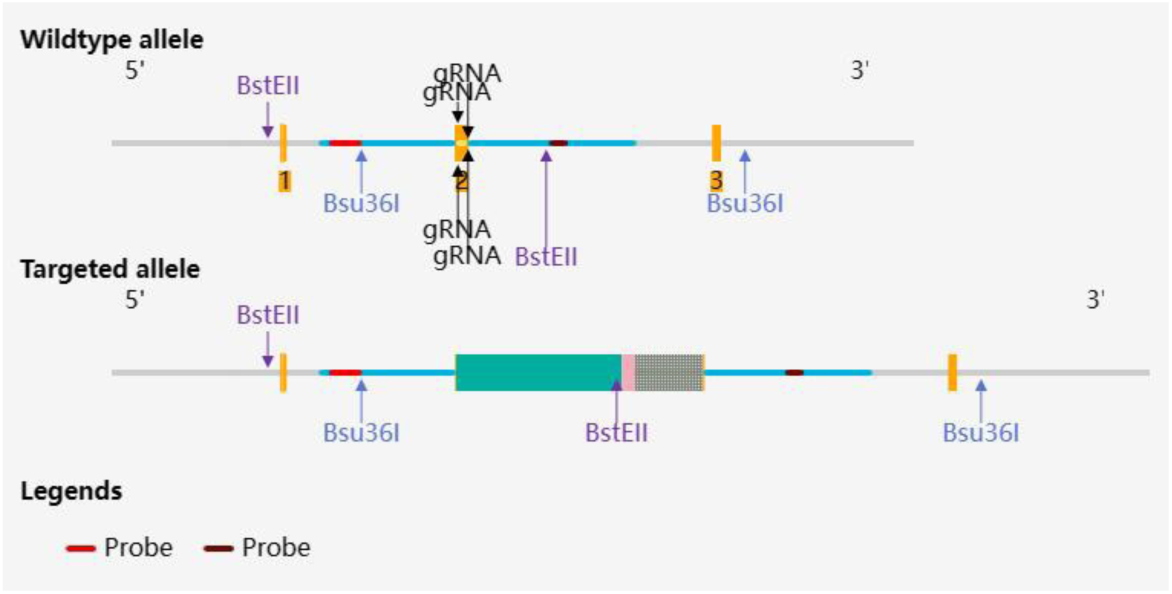

Expected Fragment Sizes for Southern Blot:

5’Probe-BstEII: 3.42 kb-WT, 4.29 kb-MT

3’Probe-Bsu36I: 4.72 kb-WT, 7.62 kb-MT

Primers for 5’Probe:

F-CTTGGGTTCGGTCCTGTACTTCTCCT

R-ACCCTTAGGATGGACAAGGAAGGA

Primers for 3’Probe:

F-CAGAGCCAAGCTGGTGTGAGTTGTAA

R-GGTTCACTTTGAGCAGTCTTCAGCTACT

All of the four rats (2, 8, 13 and 50) were confirmed correct by Southern blot analysis.

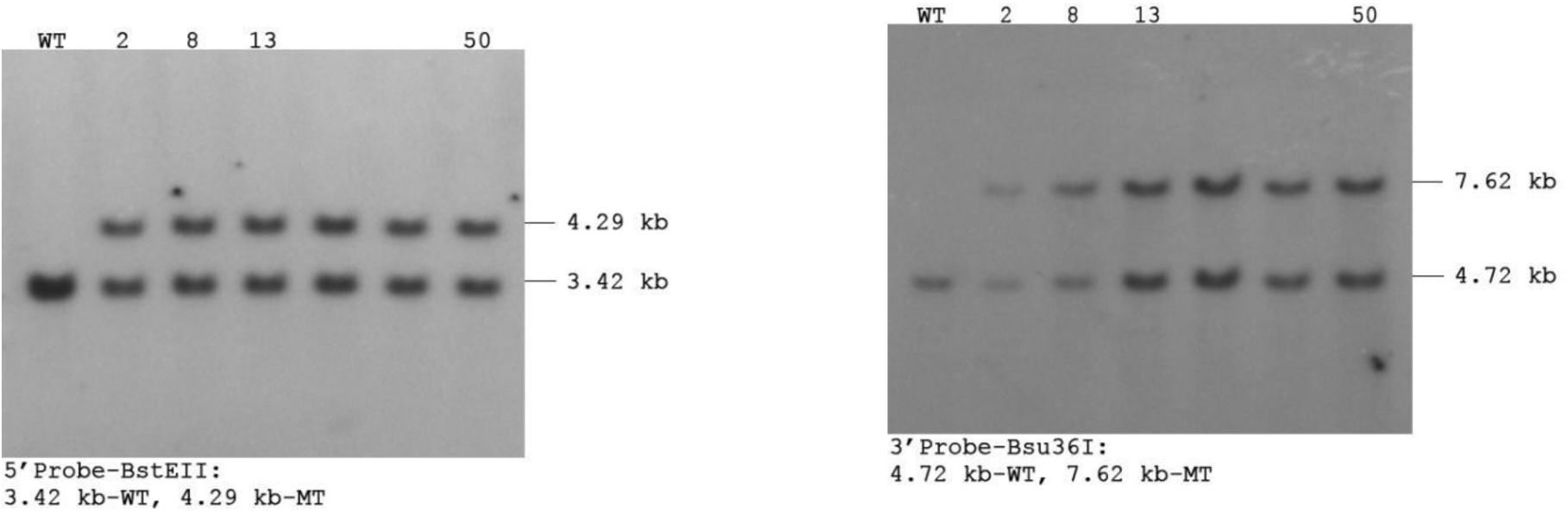

